# Islet-antigen reactive B cells display a unique phenotype and BCR repertoire in autoantibody positive and recent-onset type 1 diabetes patients

**DOI:** 10.1101/2024.06.20.599914

**Authors:** Catherine A. Nicholas, Fatima A. Tensun, Spencer A. Evans, Kevin P. Toole, Hali Broncucia, Jay R. Hesselberth, Peter A. Gottlieb, Kristen L. Wells, Mia J. Smith

**Author notes:** Corresponding authors Mia J. Smith, Kristen L. Wells. These authors contributed equally (Catherine A. Nicholas, Kristen L. Wells). Senior author (Kristen L. Wells, Mia J. Smith).

## Abstract

Autoreactive B cells play an important but ill-defined role in autoimmune type 1 diabetes (T1D). To better understand their contribution, we performed single cell gene and BCR-seq analysis on pancreatic islet antigen-reactive (IAR) B cells from the peripheral blood of nondiabetic (ND), autoantibody positive prediabetic (AAB), and recent-onset T1D individuals. We found that the frequency of IAR B cells was increased in AAB and T1D. IAR B cells from these donors had altered expression of B cell signaling, pro-inflammatory, infection, and antigen processing and presentation genes. Both AAB and T1D donors demonstrated a significant increase in certain heavy and light chain V genes, and these V genes were enriched in islet-reactivity. Public clones of IAR B cells were restricted almost entirely to AAB and T1D donors. IAR B cells were clonally expanded in the autoimmune donors, particularly the AAB group. Notably, a substantial fraction of IAR B cells in AAB and T1D donors appeared to be polyreactive, which was corroborated by analysis of recombinant monoclonal antibodies. These results expand our understanding of autoreactive B cell activation during T1D and identify unique BCR repertoire changes that may serve as biomarkers for increased disease risk.

**One Sentence Summary:** Pancreatic islet antigen-reactive B cells from individuals with prediabetes and recently diagnosed with type 1 diabetes display a unique phenotype and BCR repertoire compared to non-diabetic donors.

## INTRODUCTION

Type 1 diabetes (T1D) develops as a consequence of a sustained autoimmune attack on the insulin producing beta cells in the pancreas. T1D has historically been classified as a T cell mediated disease due to the destruction of pancreatic islet beta cells by autoreactive T cells. However, previous experiments in the non-obese diabetic (NOD) mouse model have provided evidence for autoreactive B cell involvement with disease progression. This evidence includes demonstration of their essential role in antigen presentation to T cells, protection from diabetes progression in mice lacking B cells, and requirement for islet, i.e. insulin, reactive B cells to develop autoimmune diabetes (*1-9*). Given the importance of B cells in the NOD mouse model, a phase 2 clinical trial was undertaken using the B cell depleting monoclonal antibody, Rituximab, to target CD20+ B cells in recently diagnosed individuals with T1D. The trial showed that patients treated with Rituximab have preserved beta cell function one year post-treatment (*10*, *11*). These benefits were largely lost two years after treatment when the B cell compartment had fully recovered (*12*). Despite evidence for B cell involvement in T1D, few human B cell focused studies have been completed, particularly those analyzing islet antigen-reactive (IAR) B cells. We previously analyzed insulin-binding B cells in the peripheral blood of subjects along a continuum of diabetes development and showed that anergic (unresponsive) insulin-binding B cells are lost in individuals with pre-clinical diabetes (autoantibody positive but not symptomatic) and individuals recently diagnosed with T1D (*13*, *14*). Follow-up studies in young-onset T1D revealed an increase in activated B cells within the anergic insulin-binding B cell subset, suggesting they have lost tolerance (*15*). But exactly how these B cells become activated and their role in disease progression remains unknown.

Autoantibodies produced by B cells reactive with pancreatic islet antigens, e.g. insulin (INS), glutamic acid decarboxylase 65 (GAD), insulinoma associated antigen 2 (IA2), and zinc transporter 8 (ZnT8), are found in the serum of individuals prior to onset of T1D, and are used as biomarkers to identify individuals with a high likelihood of progression to T1D (*16*, *17*). Accumulation of multiple autoantibodies in individuals with pre-clinical diabetes (prediabetes) is strongly correlated with faster progression to T1D diagnosis (*18*). Despite this, current dogma based on mouse studies suggests that autoantibodies in T1D are not pathogenic (*7*). Instead, the role of B cells in T1D is likely through (auto)antigen-presentation to T cells (*3*, *19*, *20*).

It has been shown that up to 70% of newly generated B cells in the bone marrow are self-reactive (*21*). Normally these cells are purged through central tolerance mechanisms of receptor editing or clonal deletion or by the peripheral tolerance mechanism of anergy (*22-25*). Individuals with autoimmunity, including T1D, exhibit an increase in autoreactive B cells that escape the bone marrow and enter the periphery. Importantly, these cells tend to be polyreactive, binding to two or more of the following antigens: INS, DNA, or LPS (*13*, *23*, *26*). Together these findings indicate that normal tolerance mechanisms are impaired and unregulated, autoreactive B cells play a role in the development of T1D.

Given how little is known about diabetogenic B cells, including their role in the pathogenesis of T1D and how their tolerance is broken, we sought to analyze the phenotype and BCR repertoire of islet-reactive B cells from the peripheral blood of subjects along a continuum of diabetes development. We designed a multiplexed single cell RNA sequencing (scRNA-seq) method based on LIBRA-seq (*27*) to characterize B cells reactive to three pancreatic islet antigens, INS, IA2, and GAD, as well as those reactive with the foreign antigen tetanus-toxoid (TET). While our lab has extensively studied the surface phenotype and functional properties of INS-reactive B cells using flow and mass cytometry, to our knowledge no such study has utilized scRNA-seq to study multiple IAR B cells in parallel. Our results demonstrate significant differences in the transcriptome and BCR repertoire of patients along a continuum of diabetes development compared to non-diabetic controls. These findings greatly expand our current knowledge regarding the phenotype of IAR B cells in T1D and suggest some potential mechanisms by which self-reactive B cells may lose tolerance and become activated in T1D.

## RESULTS

### B cell subsets span all stages of B cell differentiation and are present in each donor group

To more deeply phenotype IAR B cells during T1D development and obtain information regarding their BCR repertoire, we performed single cell LIBRA-(*27*), CITE-(*28*), and RNA-sequencing of INS, IA2, and GAD reactive B cells from first-degree relative non-diabetic (ND), autoantibody positive prediabetic (AAB), and recently diagnosed (<100 days) T1D donors. Peripheral blood was collected from five ND (Age: 14.2 years +/- 8.9 (mean +/- SD)), six AAB (Age: 18.2 years +/- 11.8)), and five T1D (Age: 15.8 years +/- 8.2) individuals and cryopreserved (**Table 1**). Autoantibody expression data were obtained from previous medical records and HLA testing, or HLA prediction based on mRNA expression was performed. All subjects in our study carried at least one high-risk T1D HLA allele, such as the DQ8 and/or DQ2 risk alleles (*29*). PE-labeled antigen (INS, IA2, GAD, TET) tetramers, each carrying a unique barcode sequence, were freshly made on the day of each of the scRNA-seq experiments. In addition to these antigen tetramers, we included an “Empty’’ negative control tetramer that had its four binding sites occupied by biotin without protein antigen to detect any B cells that may be reactive to streptavidin, PE, biotin, or the DNA barcode. To minimize batch effects, four PBMC samples were thawed and processed in parallel, and eight samples were processed per day.

**Table 1:**
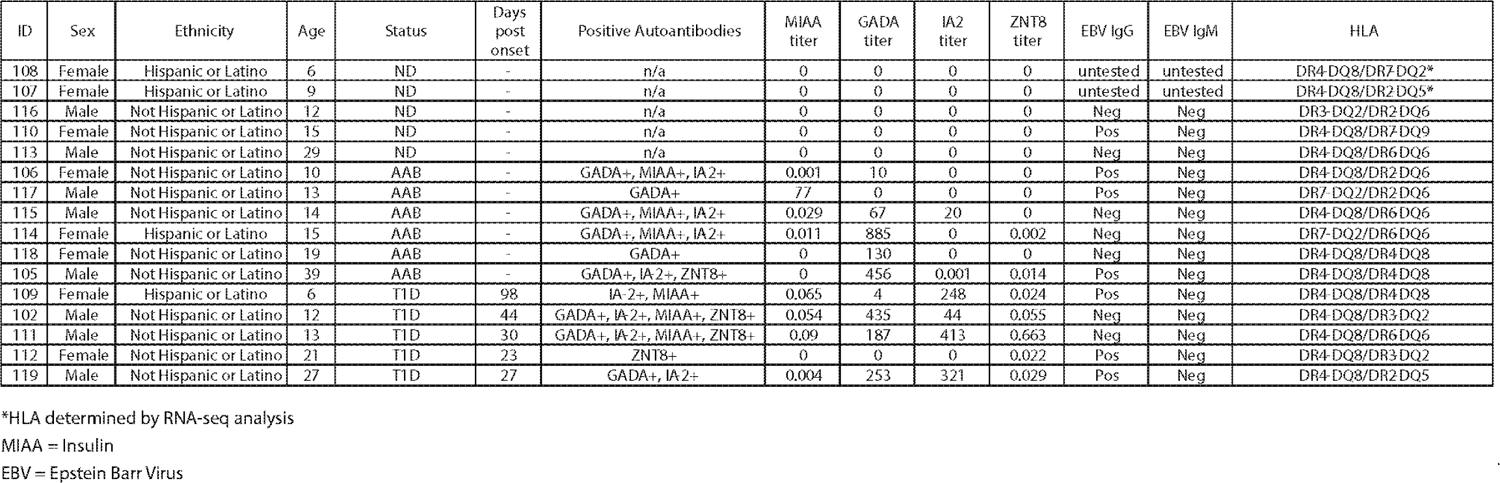
Human donor metadata. Table for 16 human donor data including autoantibodies they tested positive for, EBV antibody testing results, and HLA type.

Thawed PBMCs were incubated with the antigen and empty tetramers per LIBRA-seq (*27*), and stained with various flow and CITE-seq antibodies. PE-reactive B cells were sorted via FACS. In addition, a small number of non-PE binding B cells were also sorted and combined with the PE-binding B cells to increase the number of total cells recovered for each 10x Genomics capture (**Fig. 1A**). FACS revealed a non-significant trend toward presence of more total antigen positive binding B cells as disease progressed through stages from ND (Mean 1.46% +/- 0.25%) to AAB (Mean 1.63% +/- 0.22%) to T1D (Mean 1.99% +/- 0.36%) (**Fig. 1B**).

**Fig. 1.**
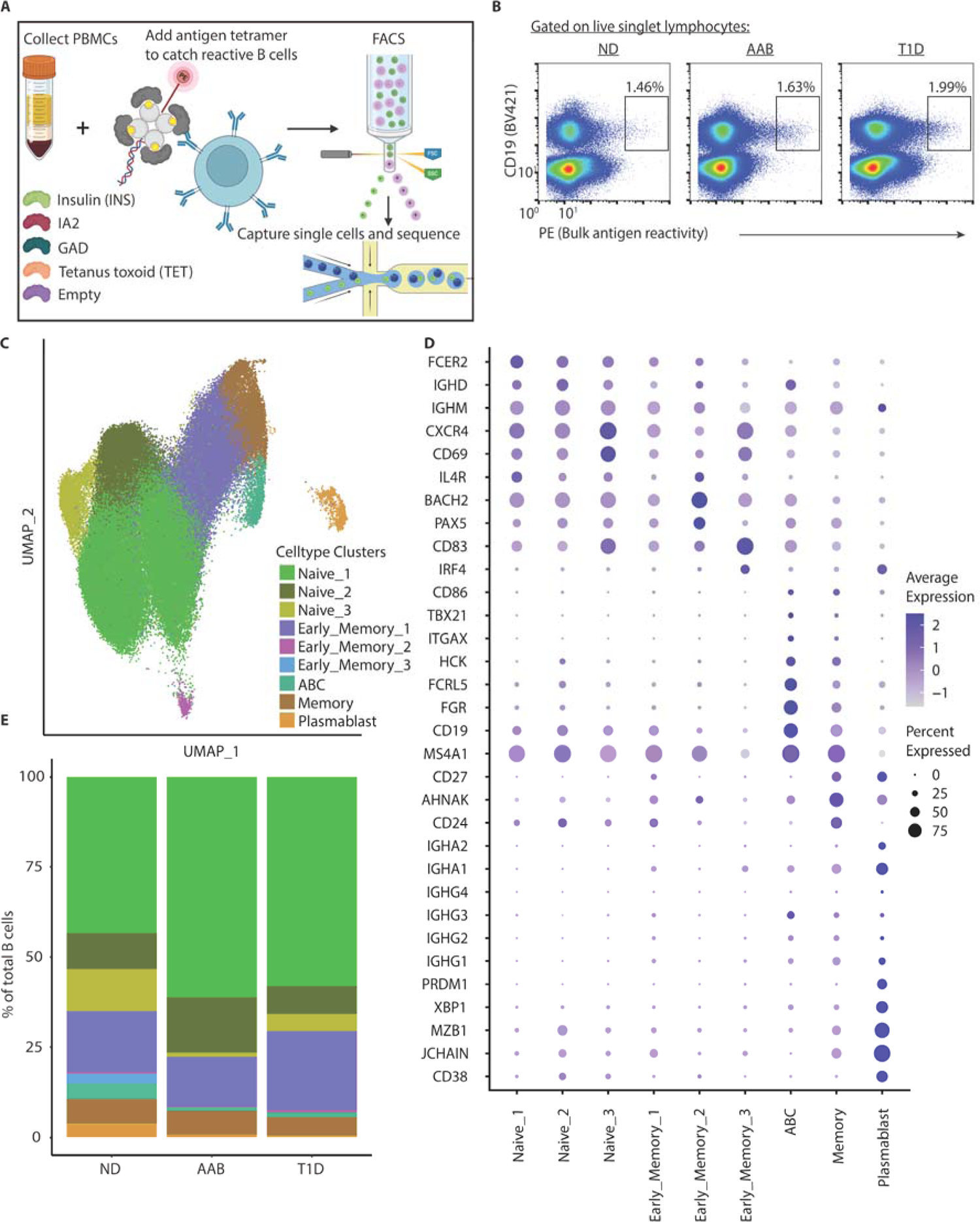
Sorted antigen-reactive B cells constitute nine B cell clusters among all donors. **(A)** Schematic overview of antigen-tetramer labeling of reactive B cells, FACS, and single cell sequencing. **(B)** Representative flow cytometry plots of how total antigen reactive (PE+) B cells from ND, AAB, and T1D donors were sorted. **(C)** UMAP plot of total B cells collected across 16 donors clustered into nine subpopulations: three naïve, three early memory, ABC, classical memory, and plasmablasts. **(D)** Dot plot of select gene expression across B cell clusters to confirm B cell subtype identity. **(E)** Stacked bar chart of B cell subtype distribution as a percent of total B cells across donor status groups.

To identify the B cell types present in our dataset irrespective of specificity, we completed standard pre-processing using the Seurat workflow (*30*) followed by sample integration and batch correction using the MNN batch correction method (**fig S1A** and **S1B**) (*31*). Following preprocessing, we identified nine unique B cell clusters in various stages of differentiation including naïve, early memory, age/autoimmune associated B cells (ABC) (*32*), classical memory, and plasmablasts (Fig. 1C and fig S2). Figure 1D confirms these cell subtypes by visualizing the expression of a selection of established genes for each cell type (*33-36*). To determine whether there were differences in representation of B cell subsets between the donor groups, we plotted the frequency of each B cell subset as a percentage of total B cells that were found in ND, AAB, and T1D subjects (**Fig. 1E**). While all B cell subsets could be found in each participant group, there was a non-significant increase in the frequency of ABC and plasmablasts in the ND group. Upon further analysis of the five ND individuals, the increase in frequency of ABC were found only in subjects 107 and 108, who, interestingly, are the ND half-sister (107) and ND fraternal twin (108) of T1D subject 109. The increase in frequency of plasmablasts was solely attributable to ND subject 110, who had a large expansion of plasmablasts among their sorted B cells. While all subjects reported that they had not been sick in the two weeks prior to their blood draw, it is possible this individual was in the early stages of an illness or had been recently vaccinated, the latter of which was not an exclusion factor for entry into our study. Thus, these five major B cell subsets were identified in our dataset and were present across ND, AAB, and T1D donor groups.

### IAR B cells are higher in frequency in AAB and T1D groups, occur in all five major B cell subsets, and tend to bind to more than one antigen

To determine the degree of IAR reactivity among the samples, we next defined a threshold of positive reactivity for each of the antigens. We calculated a binding score for each antigen using a quantile approach based on antigen-binding to non-B cells, which would reflect non-specific background binding. A binding score greater than one indicates a positive binding event and was calculated for each individual antigen. As expected, the scRNA-seq readout revealed an increase in IAR B cells as a percent of all B cells in AAB and T1D donors, though this was not significant after adjusting for multiple testing (**Fig. 2A**). Interestingly, while the proportion of INS-reactive B cells appeared stable across donor groups, we captured a greater percentage of GAD and IA2-reactive B cells from AAB and T1D compared to ND individuals. Importantly, reactivity to the empty tetramer was consistently low across all samples, indicating a low frequency of B cells reactive to any of the streptavidin, biotin, PE or DNA components of the reagent. However, we were surprised to find that the largest share of IAR B cells was composed of B cells that appeared to bind one pancreatic islet antigen and at least one additional antigen in our assay, which we termed “Islet_Multi” reactive B cells. When we analyzed the antigen reactivities found among the Islet-Multi reactive B cells, we found B cells that bound to every possible combination of the five antigen tetramers, such as some that bound both INS and GAD and others that bound all five antigen tetramers (**fig. S3**). When we quantified the percent of Islet_Multi reactive B cells that bound to each antigen tetramer, we found that Islet_Multi reactive B cells bound to the islet antigens at an increased frequency compared to the empty and TET controls (**Fig 2B**).

**Fig. 2.**
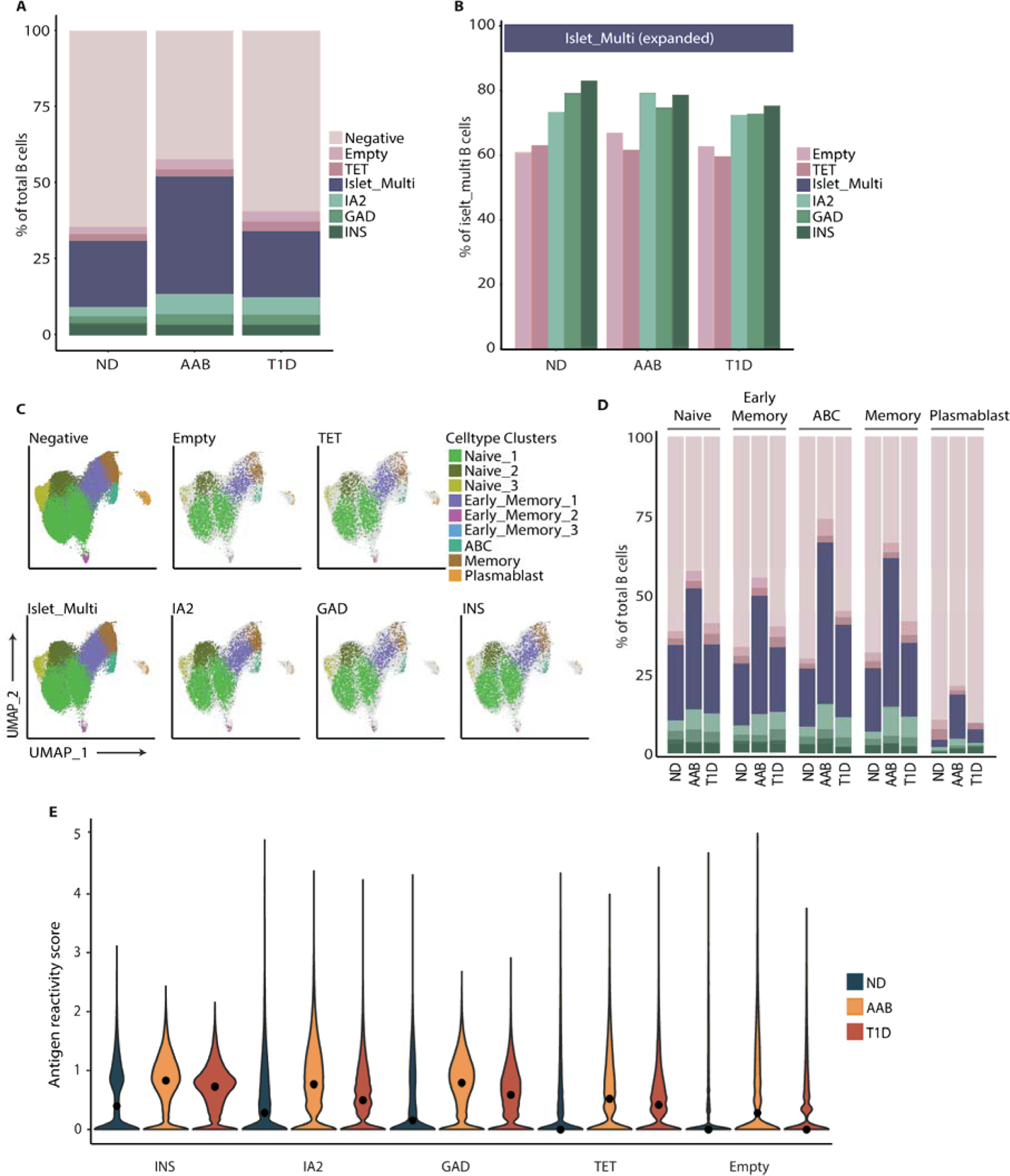
AAB and T1D donors have increased IAR B cells compared to ND donors. **(A)** Stacked bar chart of antigen reactivity as a percent of total B cells across ND, AAB, and T1D donors. **(B)** Bar chart representing the distribution of antigen reactivities found among the Islet_Multi group across donor groups. **(C)** UMAP projections of all cells colored by antigen reactivity identity **(D)** Stacked bar chart of antigen reactivity as a percent of total B cells separated by B cell subtype and donor group. **(E)** Violin plots of relative antigen reactivity score using the Quantile approach, separated by donor group.

To determine whether any of the B cell subsets (**Fig. 1**) clustered together because of shared antigen reactivity, we plotted the antigen reactive B cell groups onto the B cell subset UMAP. As seen in figure 2C, antigen-reactive B cells could be found in each of the B cell subsets, indicating B cell subsets were not clustered based on their antigen reactivity. When we quantified the frequency of antigen-reactive B cells within the five major B cell subsets in each subject group, we found that the increased proportion of IAR B cells in AAB and T1D donors is most pronounced in the more antigen experienced ABC, memory, and plasmablast B cell subsets (**Fig. 2D**). Given that IAR B cells were found at decreased but notable levels in the ND group compared to AAB and T1D donors, we next determined whether there was a difference in the level of antigen tetramer binding in ND compared to AAB and T1D donors. Most strikingly, when the antigen binding score was summarized across donor groups for each antigen, the AAB and T1D donors had higher average binding scores than ND donors (**Fig. 2E**). This indicates that the amount of antigen tetramer binding was higher in the B cells of those donors as compared to non-B cells, possibly reflecting increased affinity for antigen. The observed increased binding events for INS-reactive B cells was not due to differential insulin receptor (INSR) expression between donors (**fig. S4**). Overall, these results demonstrate that IAR B cells from AAB and T1D donors are increased in frequency, particularly in more mature B cell subsets, compared to ND donors, and tend to be polyreactive.

### IAR B cells display distinct mRNA expression patterns across T1D development

To further characterize the differences between B cells between status groups, we assessed mRNA gene expression patterns of IAR B cells. Differential gene expression between AAB and ND donors and T1D and ND donors uncovered expression changes as stages of disease progress (**Fig. 3A**). Gene set enrichment analysis identified enrichment of genes in many cell signaling pathways in AAB and T1D donors when compared to ND. We clustered a selection of KEGG pathways significantly altered in our data and grouped them into six broad signaling themes: Proinflammatory, BCR and General Signaling, Antigen Processing and Presentation, Bacterial Infection, Viral Infection, and Autoimmunity (**Fig. 3B**). As highlighted in **figure 3C**, BCR signaling pathway gene expression was significantly increased. These included increases of BLNK, PTPRC (CD45), SKAP2, CD79B, BANK1, and SYK in AAB and T1D donors - all promoting B cell activation upon antigen binding to the BCR (*37*). Downregulated genes that can contribute to inhibition of BCR signaling included LYN and the phosphatases DUSP1, DUSP2, and DUSP5 (*38*), (*39*). Additionally, IAR B cells in autoimmune donors showed increased expression of markers related to antigen presentation including CD40, CIITA, and numerous HLA genes, supporting their role as active participants in an autoreactive immune response beyond autoantibody production. The differential expression of pro-inflammatory and bacterial and viral infection pathway genes was partially overlapping, suggesting IAR B cells from AAB and T1D donors exhibit an overall pro-inflammatory phenotype compared to ND donors.

**Fig. 3.**
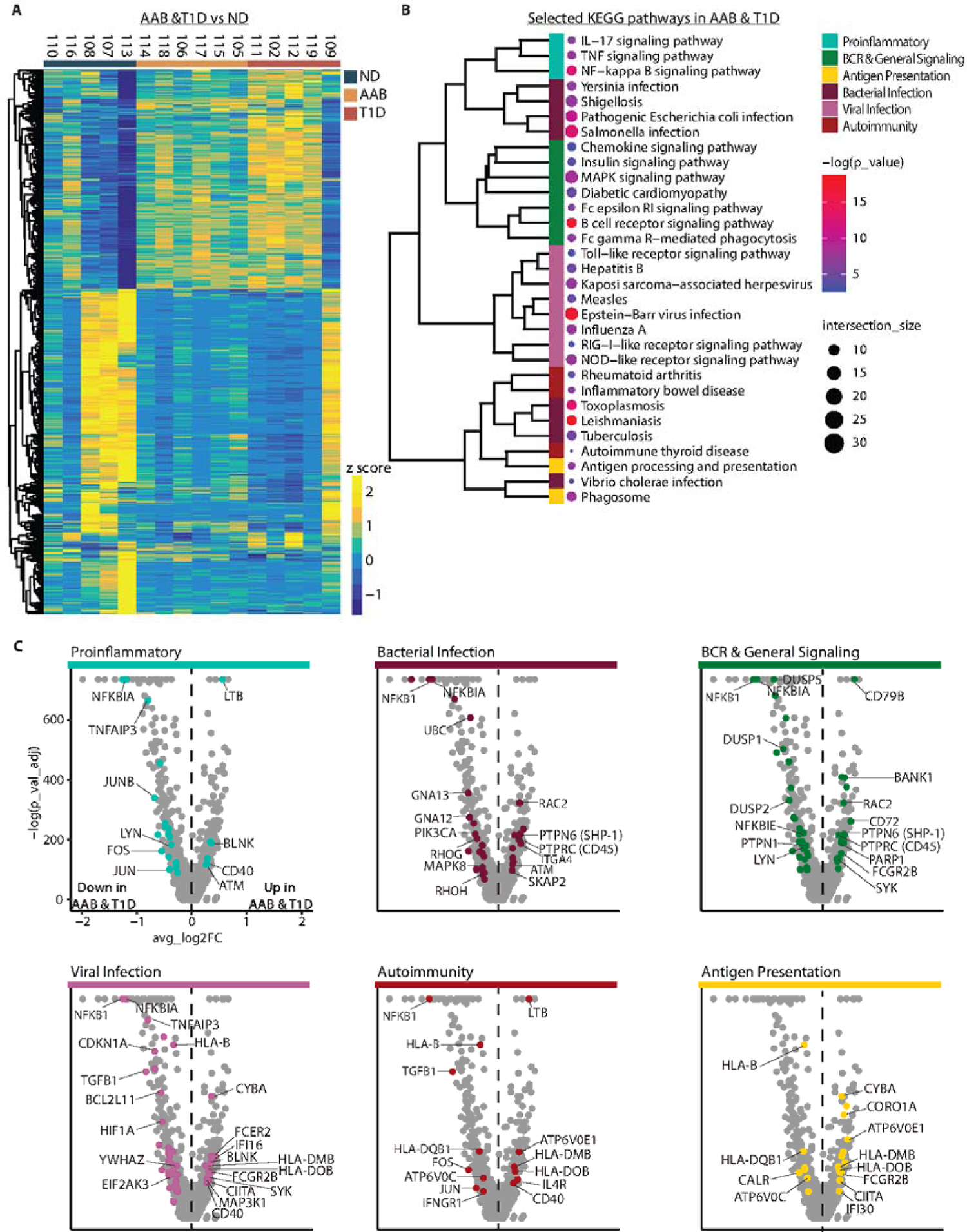
Differential mRNA gene expression patterns are found in IAR B cells in AAB and T1D donors. **(A)** Heatmap showing differentially expressed gene patterns of IAR B cells in AAB and T1D donors compared to ND donors. **(B)** Selected significantly enriched KEGG pathways for AAB and T1D donors grouped into six broad categories by general theme: Proinflammatory, BCR & General Signaling, Antigen Processing and Presentation, Bacterial Infection, Viral Infection, and Autoimmunity. Similarity between individual pathways, determined by jaccard distance, is shown by clustering dendrogram on the left side. **(C)** Volcano plots showing differentially regulated genes within the six categories defined in (B). Selected genes of interest are labeled on the plots.

Based on the differential expression analysis, donors 107, 108, and 109 appeared to have exceptionally similar gene expression patterns irrespective of disease status. These donors are siblings, with donors 108 and 109 being fraternal twins. In returning to the pre-batch corrected and normalized data, we found that those three individuals were part of an obvious batch effect when compared to all other donors (**fig. S5A**). The twins clustered together away from all other donor’s cells with their half-sister, donor 107, clustering between the twins and other donors.

Differential gene expression analysis between the twin sisters, between the three sisters, and among all donors confirmed high overlap of genes between the sisters and all donors with few genes unique to the sister comparisons (**fig. S5B**). These findings reveal that although these siblings cluster separately from the other donors, the differential gene expression patterns observed across all donors are not affected by unique gene expression due to a batch effect among these siblings.

### AAB and T1D B cells have greater BCR repertoire diversity than ND B cells

Next, we wanted to delve into the BCR repertoires of IAR B cells between donor groups. Interrogation of IAR BCR isotypes across cell types and antigen reactivities revealed that most cells in our study expressed IgM based on BCR sequencing (**Fig. 4A**). Since we sorted all PE-binding B cells, irrespective of their differentiation state (i.e. naive vs memory), these results are consistent with previous reports that ∼60-70% of peripheral blood B cells are naive and co-express IgM and IgD (*40*). It has been demonstrated previously that the CDR3 regions of autoreactive BCRs tend to be longer and contain more positively charged amino acids (*41*, *42*). We found a modest increase in CDR3 amino acid length and positive charges in the IAR B cells (**Fig. 4B** and **Fig. 4C**). There was no difference in level of somatic hypermutation (SHM) in the heavy or light chains of IAR B cells among donors, likely due to the high level of IgM isotype of B cells in the samples (**Fig. 4D** and **Fig. 4E**). However, when we calculated a Shannon diversity score across cell types and donor groups in total B cells, we discovered significantly higher diversity in the autoimmune AAB and T1D donors compared to ND individuals (**Fig. 4F**). This trend was largely maintained in more mature cell populations of ABC, memory, and plasmablasts. The increased diversity of BCRs among autoimmune donors may reflect defects in central and peripheral B cell tolerance in individuals with T1D (*23*, *26*), allowing more autoreactive B cells to escape into the periphery.

**Fig. 4.**
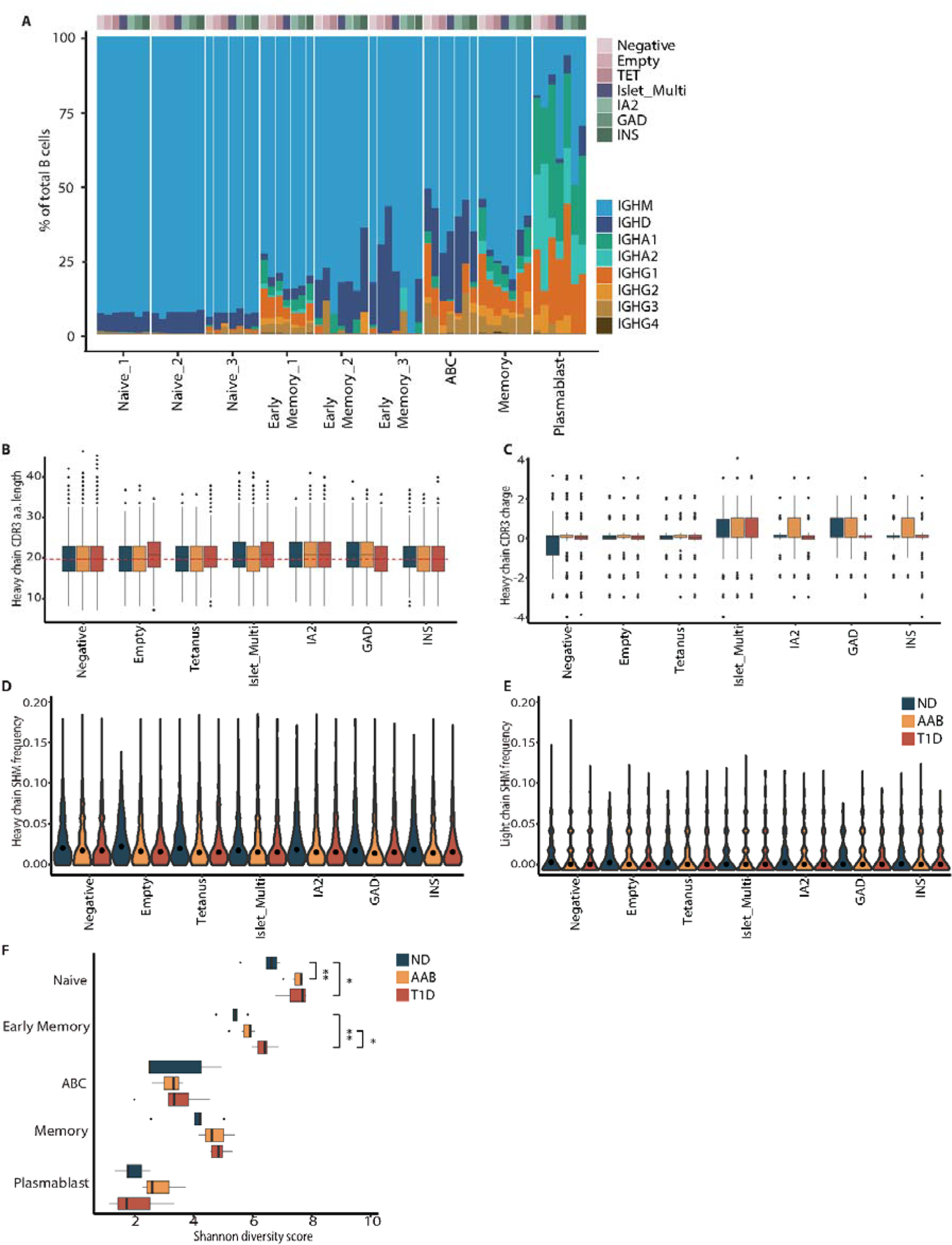
B cell receptor analysis reveals increased diversity in IAR B cells of AAB and T1D donors. **(A)** Stacked bar plot of BCR isotype usage determined by V(D)J analysis as a percent of all B cells. Cells are divided by subtype and antigen reactivity. **(B)** Amino acid length of heavy chain CDR3s for each antigen and donor group. **(C)** Heavy chain CDR3 net charge for each antigen and donor group. **(D** and **E)** Violin plots of somatic hypermutation (SHM) frequency for each antigen and donor group for the heavy chain (D) and light chain (E). **(F)** Shannon diversity score for each B cell subtype within each donor group. A t-test was calculated for each donor group comparison. Significant differences are shown with * p < 0.05 and ** p < 0.01.

### AAB and T1D donors exhibit a unique BCR repertoire with significant enrichment of islet-reactive heavy and light chain gene pairs

To determine whether there is any T1D-associated BCR gene usage, we next analyzed the overall heavy and light chain V gene repertoire across ND, AAB, and T1D B cells, irrespective of their antigen specificity (**Fig. 5A, fig S6A**). We found significant differences in the frequency of heavy and light V genes identified in T1D and AAB individuals compared to ND individuals. An odds ratio test revealed IGHV4-4, IGHV1-3, IGHV3-64, IGHV3-66, IGHV4-61, and IGHV1-18 gene usage was significantly greater in AAB and T1D donors than in ND (**Fig. 5B**). Light chain V genes used more frequently in both AAB and T1D were IGLV9-49 and IGKV1-8 (**fig S6B**). Conversely, several heavy and light chain genes were expressed less among AAB and T1D donors; these included IGHV3-30, IGHV1-2, IGHV1-69D, IGHV3-11, IGHV4-30-4, IGHV2-70D, IGHV1-69, IGLV2-23, IGLV3-1, IGLV3-19, IGLV2-14, and IGKV1-33 (Fig. 5B and fig S6B).

**Fig. 5.**
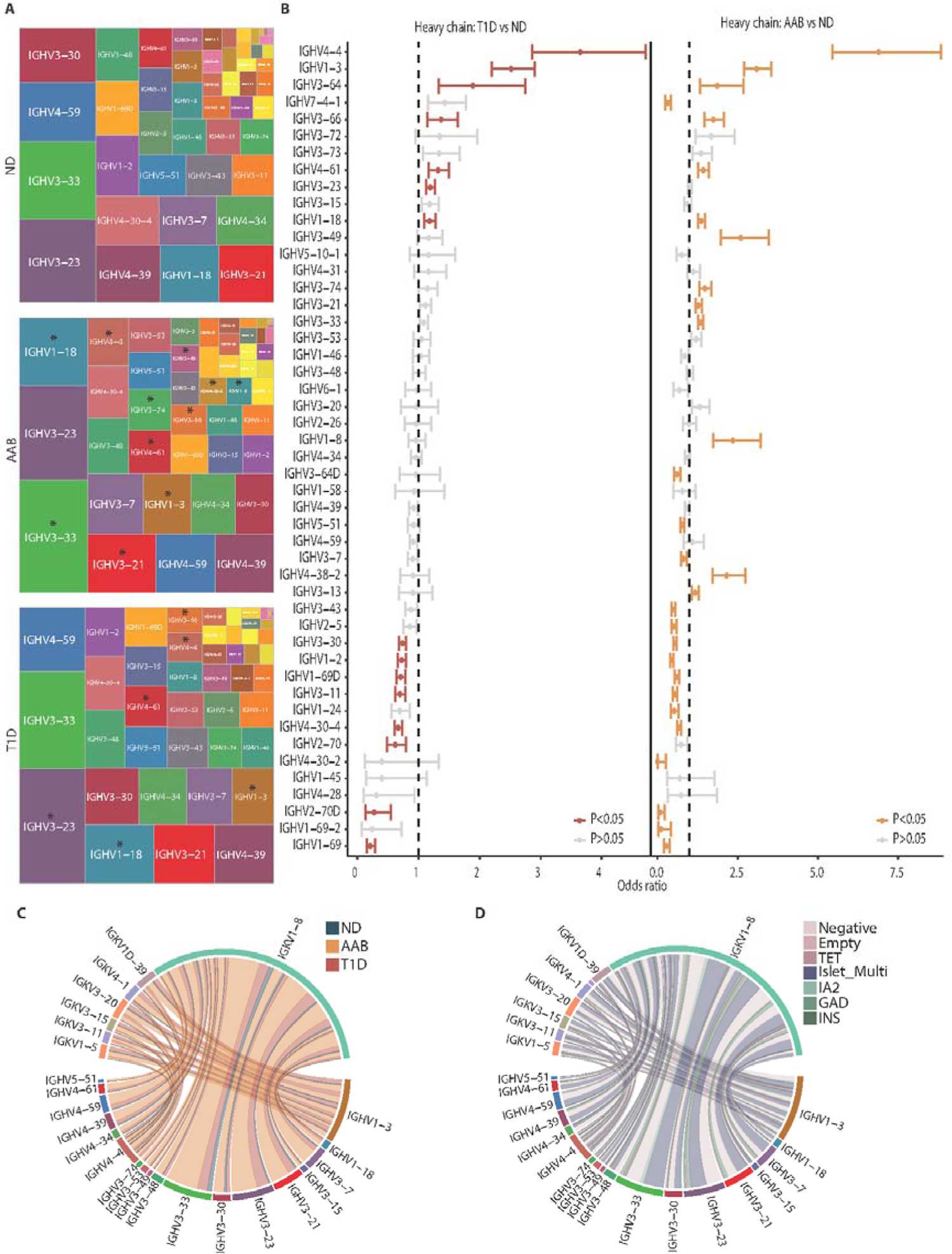
AAB and T1D donors have significant enrichment for particular heavy chain V gene usages. **(A)** Total B cells for each donor group were assessed for heavy chain V gene usage and plotted such that the size of each box is proportional to the frequency with which that gene was found among donors. Asterisks mark genes that were found to be significantly upregulated in AAB and T1D donors compared to ND. **(B)** Odds ratios were performed for heavy chain V gene usage between T1D and ND donors and AAB and ND donors. P value is indicated by color where red indicates p < 0.05 for T1D vs ND and orange indicates p < 0.05 for AAB vs ND. **(C)** Circos plot of significantly enriched heavy and light chain gene pairings in T1D and AAB vs ND. Heavy and light pairs are connected by lines colored by donor group and are thickened by the number of cells that used each segment. **(D)** As in (C), but pairs are colored by antigen reactivity.

We next determined whether expression of any unique heavy and light chain gene pairings was increased in the autoimmune donors compared to ND donors. Multiple heavy and light chain gene pairs found included pairs that used the significantly increased heavy chain genes IGHV4-4, IGHV1-3, IGHV4-61, and IGHV1-18, as well as the light chain gene IGKV1-8 (**Fig. 5C**). Interestingly, most of these pairings were observed in the AAB group. When we analyzed antigen reactivity of these paired heavy and light chains, we found that many bound islet antigens and were among the Islet_Multi group (**Fig. 5D**). Altogether, we found unique BCR repertoire features in AAB and T1D donors compared to ND. These included significant increases in heavy-light chain gene pairs that exhibit islet-reactivity.

### Clonally expanded IAR B cells are predominantly found among AAB and T1D donors

We next examined whether any IAR B cells were clonally expanded in the AAB and/or T1D donors compared to ND donors. While there was relatively equal representation of Negative, Empty, and TET reactive clones in ND, AAB, and T1D donors, we found that clonally expanded IAR B cells were almost exclusively found among the autoimmune donors, particularly in the AAB group (**Fig. 6A** and **6B, Supplementary File 1**). Given that previous studies have identified public (i.e. common) TCRs that associate with T1D disease (*43*, *44*), we searched for BCR sequences in our dataset that were enriched across donors. We identified 89 public B cell clonal sequences across 16 donors with diverse V gene pairings (**Fig. 6C**). These public sequences varied in their representation among individuals and their donor group status, but consisted of predominantly autoimmune AAB and T1D donors (**Fig. 6D**). When we analyzed the antigen reactivity of these public BCR sequences, we found that many were islet antigen-reactive (**Supplementary File 1).** Overall, these findings demonstrate that clonal expansions of both private and public IAR BCR sequences occur and are found predominantly in AAB and T1D donors.

**Fig. 6.**
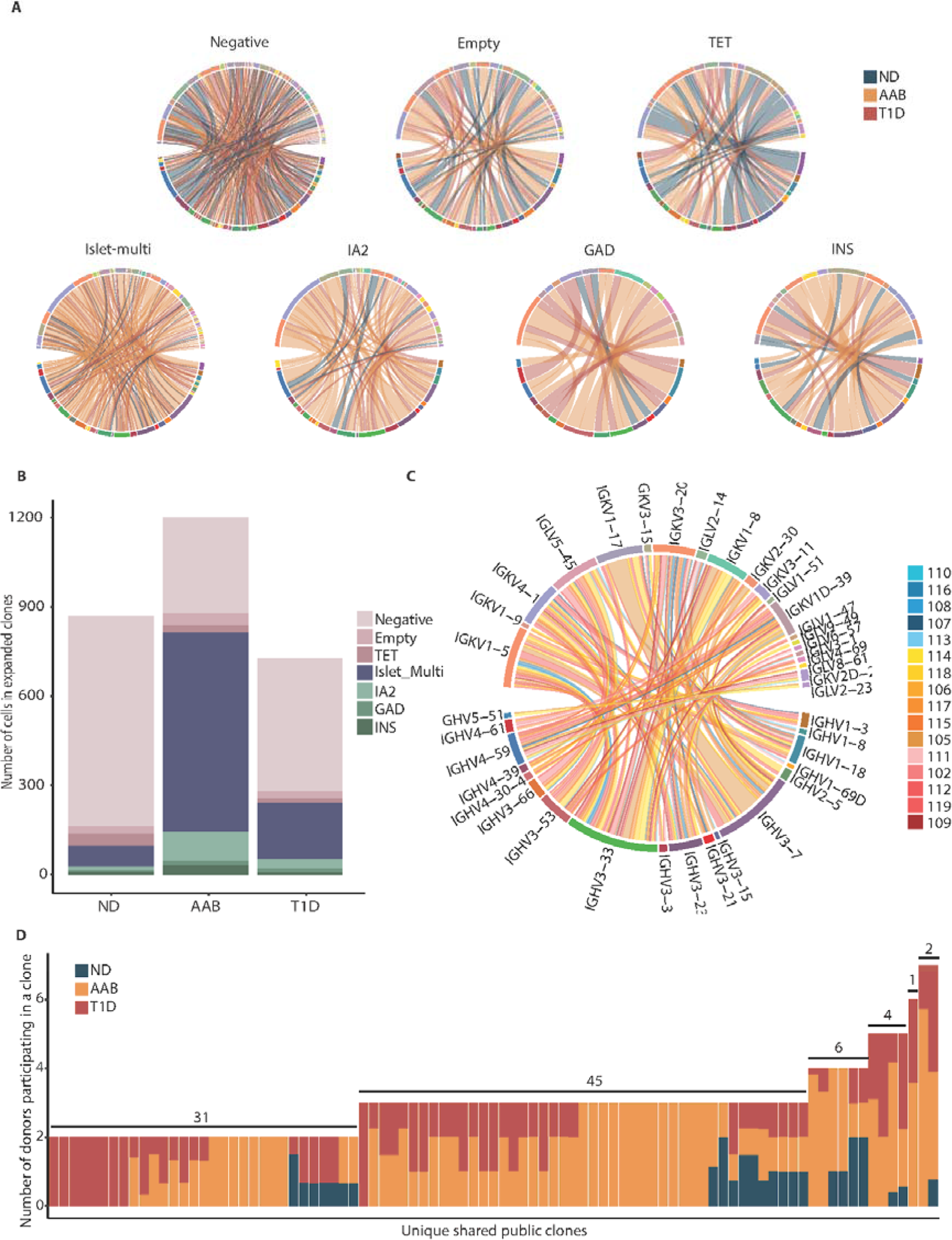
Clonal expansions of IAR B cells are increased in AAB and T1D donors. **(A)** Circos plots of heavy and light V gene pairings from private (within a single individual donor) clonally expanded cells. Segment connections are colored by donor status groups and plots are separated by antigen reactivity. **(B)** Stacked bar plot of the absolute number of private clonally expanded cells grouped by donor type and colored by antigen reactivity. **(C)** Circos plot of V gene pairs from public (shared among at least two donors) clonally expanded cells. Segment connections are colored by individual donor ID. Shades of blue represent ND, shades of yellow represent AAB, and shades of red represent T1D donors. **(D)** Stacked bar plot with the number of individual donors participating in each shared public clone (n=89) colored by donor status.

### Recombinant antibodies validate the antigen-reactivity of BCR sequences

To validate the antigen reactivity of the BCR sequences identified using our quantile approach, we selected 27 paired heavy and light chain V(D)J sequences (**Supplementary File 2**) for recombinant antibody production. Sequences were selected from the private clones in AAB and T1D donors that were found to bind each of the islet antigens, tetanus, and some that were in the Islet_Multi group (**Table 2**). As shown in Figure 7, results demonstrated that the recombinant antibodies were positive for each of the antigens implicated in cell binding studies, although some only became positive at the highest concentration of antibody added (**Fig. 7A-D**). These results indicate that while the antigen reactivities confirmed our quantile scoring method, many of these antibodies were low affinity for their antigen. This is not surprising given all but two were IgM isotype and had not undergone extensive somatic hypermutations. We next tested the reactivity of the antibodies that were identified to have bound to two or more antigens in the Islet-Multi group against each of the islet antigens, tetanus, DNA, and PE. Four of the nine Islet-Multi sequences tested were from IgG class switched cells. We found that 7/9 B cells identified as Islet-Multi were indeed polyreactive even at lower concentrations, 1/9 was polyreactive only at the highest concentration, and 1/9 was PE-reactive (**Fig. 7E**). Finally, while our quantile scoring approach closely agreed with the previous method (LIBRA-seq) to identify antigen-reactive B cells via scRNA-seq of single-reactive B cells, we found that the two methods did not agree well for cells that were called Islet_Multi (**fig. S7**). Since each of the tested Islet_Multi antibodies were reactive to at least one antigen, and most were confirmed to be polyreactive (**Fig. 7E**), we found that our quantile scoring method was much more accurate than the LIBRA-seq method for identifying polyreactive B cells.

**Fig. 7.**
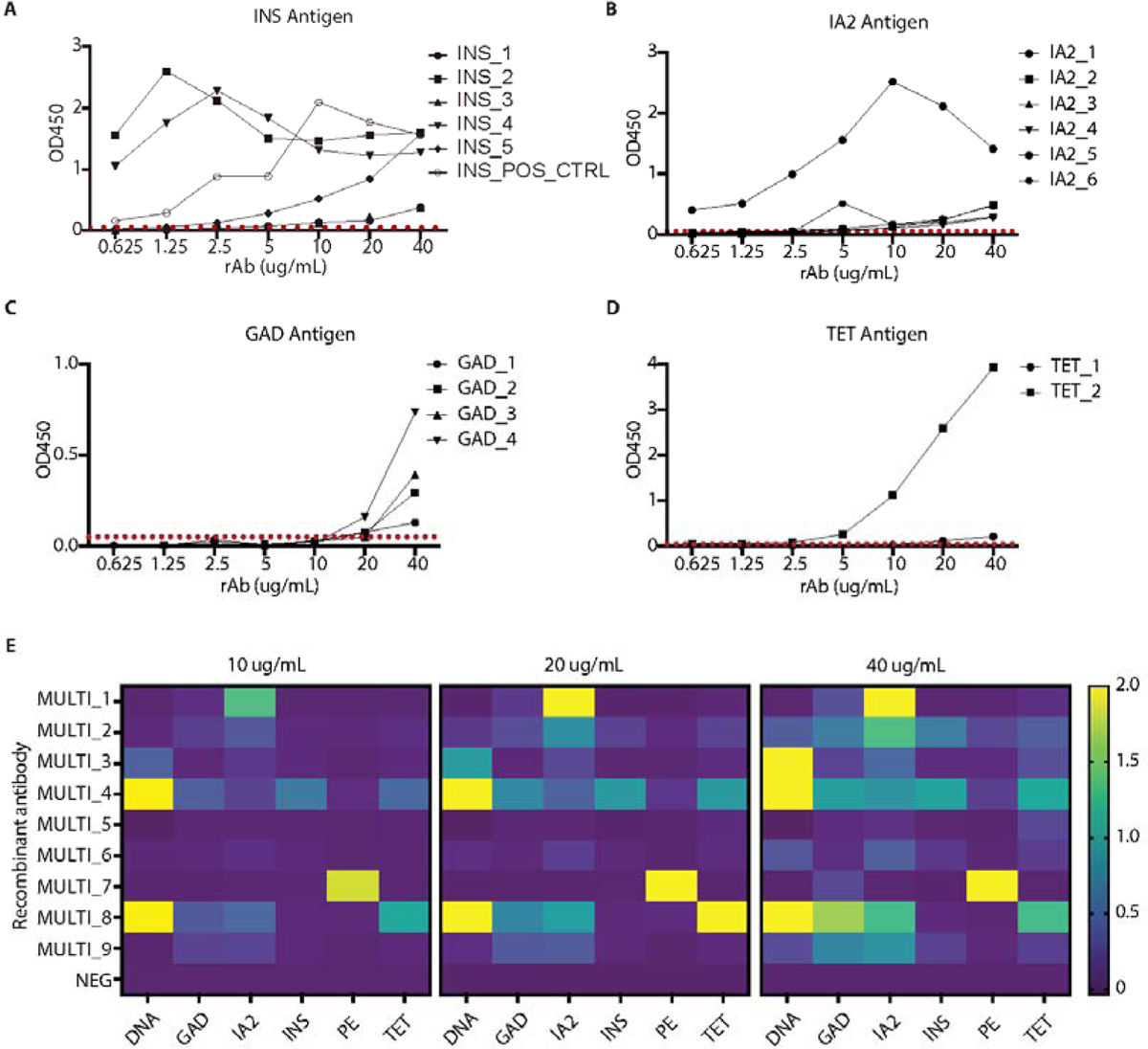
Antigen reactive BCR V(D)J sequences were confirmed by ELISA. **(A-D)** Sequences identified from IAR B cells in AAB and T1D donors were used to make recombinant monoclonal antibodies, which were tested by ELISA against their identified antigen. One positive control INS reactive sequence was tested alongside our other experimentally indicated INS reactive sequences. Plots show OD450 values of three technical replicates at each antibody concentration. A red dashed line indicates the level of assay background for each antigen tested. (E) Nine sequences identified from Multi-islet antigen reactive B cells in AAB and T1D donors were used to make recombinant antibodies, which were tested by ELISA against all experimental antigens present, including PE and DNA that were present in the tetramer probes. A negative control antibody is included for comparison. Heatmap representing OD450 values for each antibody’s reactivity to each given antigen. OD450 values ≥2.0 are yellow.

**Table 2:**
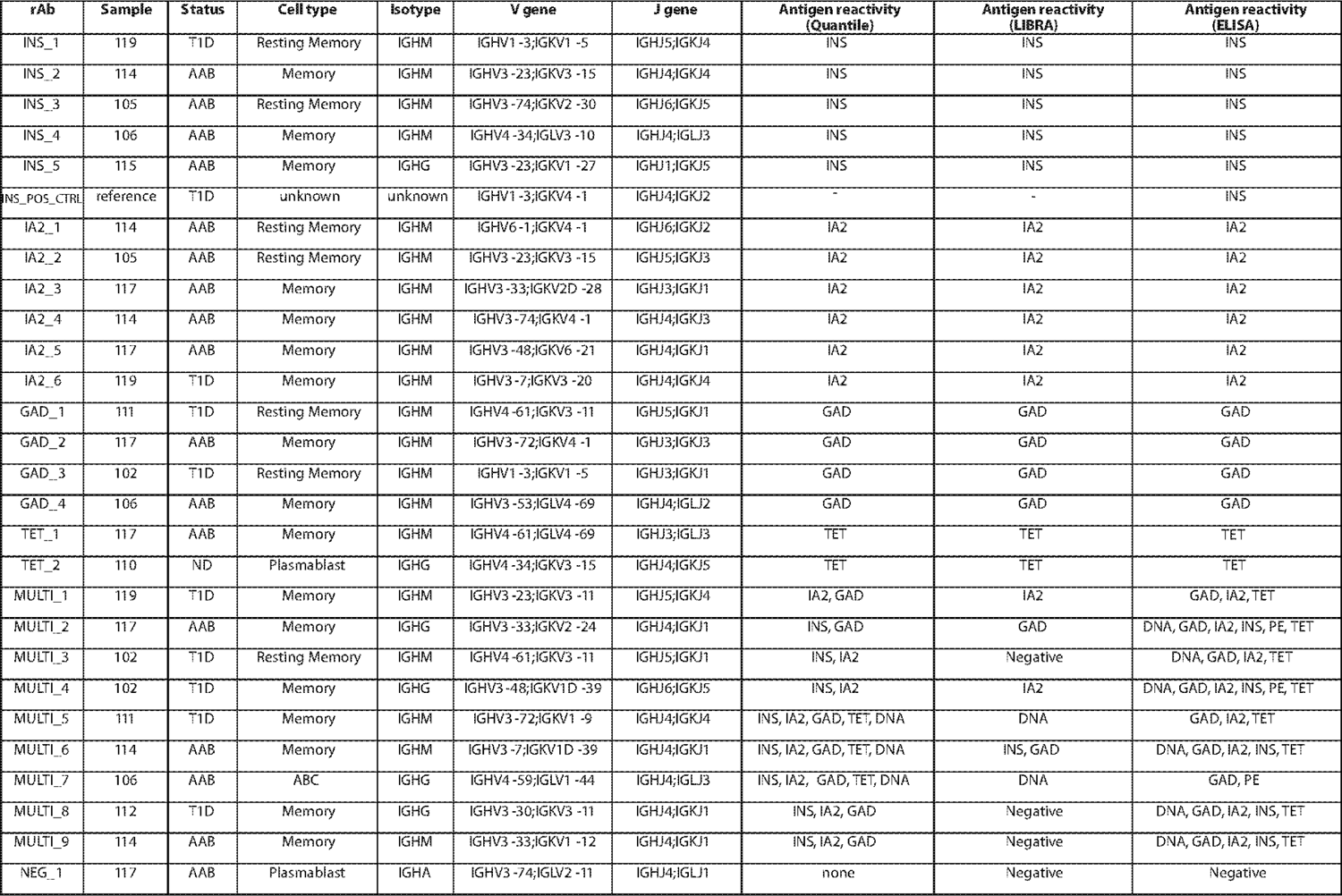
Recombinant antibody gene information and results from quantile, Libra, and **ELISA testing.** For each recombinant antibody made, the information about which sample and cell type it came from, V(D)J gene data, and the results from each of the tests is included. Full sequence information is found in supplementary file 2.

## DISCUSSION

Although most studies of T1D have focused on T cells, B cells are essential to the pathogenesis of T1D. Here we performed the first scRNA-seq study that focuses on the phenotypes and BCR repertoire of peripheral blood B cells in both AAB and T1D donors. Moreover, by studying INS, GAD, and IA-2 reactive B cells, we incorporated analyses of the pertinent antigen-reactive B cells thought to contribute to the pathogenesis of T1D. Our findings revealed that IAR B cells from AAB and recent onset T1D donors display a unique phenotype including altered gene expression in pathways related to cell signaling, antigen presentation, and inflammation compared to ND donors. In addition, IAR B cells from AAB and T1D donors are clonally expanded, tend to be polyreactive, and exhibit a distinct BCR repertoire compared to ND donors. These findings provide an excellent backdrop and resource to pursue further studies on the unique features of IAR B cells during progression to T1D. They will additionally aid in development of more tailored B cell targeted therapies and to monitor B cell response to therapy.

Our analysis of AAB patients begins to unlock early disease phenotypes specific to IAR B cells. AAB individuals have phenotypic patterns similar to T1D donors, yet remain distinct with increased unique V(D)J gene usage over ND donors and greater representation of clonally expanded IAR B cells compared to T1D donors. These findings suggest that altered B cell phenotypes may be most pronounced prior to clinical diagnosis of T1D. By the time of clinical diagnosis, B cells are likely less actively participating in direct autoimmune attack, and therefore, the unique gene signatures that we identified may be waning. Future studies will be aimed at extending these analyses to more ND autoantibody negative first-degree relatives to determine whether a B cell gene signature exists prior to seroconversion, and therefore could be used as the earliest biomarkers of disease described to date.

During our investigation into differentially expressed genes in IAR B cells in AAB and T1D donors compared to ND donors, we found several disease-related markers. Significant differences in mRNA encoding BCR signaling proteins including *BLNK, LTB, CD40, PTPN6 (SHP-1), LYN, CD79B, BANK1*, and *SYK* were observed in AAB and T1D samples compared to ND donors. Many of these genes function as activating or inhibiting signals upon antigen binding to the BCR, depending on the context. LYN, a Src tyrosine kinase known for dual roles in BCR signaling including both activation and inhibition was downregulated in AAB and T1D donors. Studies in mice have demonstrated that the inhibitory role of *LYN* is important in maintaining B cell tolerance to self-antigens (*45*, *46*). Perhaps the decreased expression of *LYN* in AAB and T1D donors contributes to the rogue activation of IAR B cells. Our data also revealed that IAR B cells may function as antigen presenting cells, as evidenced by increased expression of genes related to this pathway in AAB and T1D donors. While mouse studies have suggested the role of B cells in T1D involves antigen presentation to T cells (*3*, *19*, *20*), this has never been formally demonstrated in humans. Differential gene expression showed IAR B cells from AAB and T1D donors are enriched in genes related to inflammation, activation, and infection. Environmental triggers, such as viral infection, have been proposed as precipitating events during the pathogenesis of T1D (*47*), (*48*). While none of our subjects reported illness at the time of blood draw, infection can lead to loss of B cell tolerance, activation, and autoantibody production (*49*, *50*, *51*). Given that Epstein Barr virus (EBV) infects B cells, we tested 14/16 donors for evidence of recent or prior infection with EBV. We found that 3/5 T1D, 3/6 AAB, and 1/3 ND donors were positive for anti-EBV IgG (**Table 1**). Taken together, IAR B cells in AAB and T1D donors display signs of activation and loss of B cell anergy, which is consistent with previous reports (*13*, *15*).

Intriguingly, we detected a substantial number of polyreactive IAR B cells in our samples. Polyreactivity among early transitional and mature B cells has previously been established as a feature of multiple autoimmune diseases, including T1D (*42*, *52*). Strikingly, the AAB donors in our study had the most Islet_Multi polyreactive expanded clones, even more than T1D donors. We hypothesize that these findings reflect increased immune activation in AAB donors during the earlier stages of disease. The increase in polyreactivity and clonal expansion of IAR B cells may indicate B cells could be initially activated by an antigen other than islet antigens, such as DNA, or potentially by a foreign antigen, such as a virus. Then, through molecular mimicry and/or affinity maturation towards islet reactivity, those B cells begin targeting islet antigens (*53-56*). Future studies in the lab will address this possibility in greater detail. Furthermore, it is important to note that these polyreactive B cells were identified using our quantile scoring approach and not via the previously published LIBRA-seq approach (*27*).

The LIBRA-seq method combines CLR normalization with a z-score to define a cell as antigen binding. This method is consequently biased against the possibility of finding cells reactive to more than a single antigen. This is different from our quantile method where we determined a cutoff for antigen reactivity on a per-sample basis using the non-B cells captured in the experiment to set a threshold for positivity, which was better able to identify the polyreactive B cells.

Even though all donors in this study shared HLA T1D risk alleles (*29*) and all ND donors were first-degree relatives of someone with T1D, we discovered BCR repertoire differences between our donor groups. In particular, IGHV4-4, IGHV1-3, and IGHV3-64 were significantly enriched in individuals experiencing islet autoimmunity compared to ND donors. Interestingly, IGHV1-3 has previously been shown to be reactive to the EBV protein LMP-1 (*57*). In addition, IGHV4-4 gene usage was enriched in antibodies found to target both rheumatoid arthritis (RA) associated antigens and those from bacterial species (*58*). These results may support an infectious environmental instigator of T1D hypothesis. Interestingly, AAB and T1D donor B cells showed a significantly decreased usage of IGHV1-69 which has previously been enriched in RA and multiple sclerosis (MS) (*59-61*). IGHV1-69 polymorphisms have also been linked with varying responses to both natural infection and vaccine response (*62*). Despite the fact that IGHV4-34 gene usage is over-represented in many autoimmune diseases, including systemic lupus erythematosus (SLE), MS, and RA (*41*, *63*), we did not find it to be enriched in our AAB and T1D donors. However, we did find significant enrichment for light chain IGKV1-8 in AAB and T1D donors compared to ND, and this paired with IGHV4-34 was found to be represented at a greater frequency in the autoimmune samples in this study. Moreover, there is 98.38% sequence similarity between IGHV4-4 and IGHV4-34 which may warrant further exploration.

Taken together, our findings revealed significant differences in the BCR repertoire of AAB and T1D donors compared to ND. These results represent a foundation from which to consider new therapeutics to target only the pathogenic B cells, while preserving non-pathogenic B cells, as well as the possibility of using the BCR repertoire as a biomarker of increased risk for disease development and/or to monitor response to therapy.

While this study has expanded our understanding of the role of B cells and B cell phenotypes during development of T1D, it has a few limitations. We sequenced peripheral blood B cells from 5-6 donors per group, which is a relatively small number of subjects. Despite this, we found many significant differences in gene expression and BCR repertoire between donor groups, and adding more individuals would further improve our statistical power. In addition, by sequencing more individuals in the future we hope to compare B cell phenotypes of young (<12 years) to adult (>18 years) AAB and T1D donors, given previous studies have demonstrated B cells likely play a more important role in individuals diagnosed with T1D at a young age (*64-66*). By focusing this study on peripheral blood B cells, we may have identified phenotypes that are not reflective of pathogenic B cells localized to the target tissue, such as the pancreas or pancreatic lymph nodes. Studies in our laboratory are currently underway to address this possibility. Finally, a large portion of the B cells that we captured were naive or IgM-expressing B cells. Recombinant monoclonal antibodies demonstrated that the affinities of the B cells were largely low affinity, which is not unexpected given the low level of somatic hypermutations found in naive, non-class switched B cells. However, autoreactive IgM+ cells have been found to still contribute to autoimmune pathogenesis in RA, autoimmune hemolytic anemia, and pemphigus or autoimmune neuropathy (*67*). Future studies in the lab also aim to sequence more memory-like B cells to further characterize these B cell subsets and their differences in T1D vs ND donors.

In summary, this study revealed significant differences in gene expression and the BCR repertoire of peripheral blood IAR B cells in individuals with prediabetes and T1D compared to ND donors. These findings provide insight into how autoreactive B cells can be solicited to participate in an autoimmune response, demonstrate the breath of polyreactivity that is present among self-reactive B cells, and suggest pathways and/or targets that could be manipulated for future therapeutics to treat or prevent T1D.

## MATERIALS AND METHODS

### Study Design

This study sought to determine whether differences in the transcriptional phenotype and BCR repertoire of IAR B cells exist during development of autoimmunity. We generated uniquely barcoded PE-labeled antigen tetramers for INS, GAD, IA-2, and TET to identify antigen-reactive B cells using scRNA-seq. PBMCs from five ND first-degree relatives, six AAB subjects, and five recent-onset T1D patients were incubated with the various tetramers. Then, PE-binding B cells were sorted and loaded onto the 10X Genomics platform to be analyzed for gene expression and BCR repertoire differences. Recombinant antibodies were generated to confirm antigen-reactivity of some of the identified BCR sequences.

### Study participants

All participants were enrolled under study protocols approved by the Colorado Multiple Institutional Review Board (COMIRB #01-384). Written informed consent was obtained from each adult participant, or from parents or guardians of participants less than 18 years old. Assent was obtained from participants over the age of 7 years old who were cognitively able to consent. Participants were compensated for completing study procedures. All procedures were performed in accordance with COMIRB guidelines and regulations. Demographics and other data about research participants can be found in Table 1. Participants were recruited at the Barbara Davis Center at the University of Colorado Anschutz Medical Campus. Eligible subjects were male or female who had been recently diagnosed (<100 days) with T1D, Stage 1 or Stage 2 autoantibody positive prediabetic, or first-degree relatives of someone diagnosed with T1D. Inclusion criteria for the first-degree relative group included: 1) autoantibody negative and 2) no history of autoimmune disease. All participants were required to have no history of illness within the last two weeks. HLA haplotypes and autoantibody levels against INS, IA-2, GAD65, and ZnT8 were determined at the CLIA certified Autoantibody/HLA Service Center at the Barbara Davis Center for Diabetes.

### Blood sample processing

Between 15 – 45 mL of blood was drawn into sodium heparin tubes. The blood was rocked for no more than 2 hours at room temperature prior to PBMC isolation using Ficoll density centrifugation. DNA for HLA typing was extracted from granulocytes using the DNA midi kit (Qiagen, Cat#51185). Diluted (1:1) plasma was collected from the Ficoll layer and frozen. PBMCs were cryo-frozen prior to use in the described experiments.

### IA2 protein expression

A gene block was ordered from IDT containing the reverse transcription DNA sequence of most likely codons taken from the amino acid sequence of residues 605-979 of IA2 (i.e. PTPRN, which is known to bind IA2 specific autoantibodies in T1D individuals) and the appropriate flanking sequences for expression in our system (Supplementary File 3). 100 ng of the IA2 gene block was combined with PDonor 221 plasmid, BP clonase (Invitrogen, Cat#56481), and DH5-alpha competent cells were transformed and grown. Next, the IA2 transformed plasmid DNA was isolated and combined with Pet53dest plasmid, LR clonase (Invitrogen, Cat#56484), and DH5-alpha competent cells were transformed and grown. After this, the IA2 geneblock within the Pet53dest plasmid was isolated and mixed with BL21DE3 competent cells and grown prior to IPTG (Millipore Sigma, Cat#367-93-1) and EtOH induction of gene expression. The His-tagged IA2 was then purified with a NiNTA Agarose bead (Qiagen, Cat#1018244) packed column fraction collection. Pure IA2 fractions were determined by gel electrophoresis and proper folding and IA2 epitope accessibility was confirmed by the Barbara Davis Center Autoantibody Core. All cells and materials used in IA2 production were donated by Jay Hesselberth.

### Generation of antigen tetramers

Human recombinant insulin (Millipore Sigma, Cat#19278), GAD65 (Diamyd Medical, Cat#45-08029-01), IA2 (described above), and tetanus-toxoid (Colorado Serum Company) were biotinylated using the Thermo EZ-Link Sulfo-NHS-LC-Biotin Kit (Cat#A39257) with a 20 molar ratio of biotin to protein. Excess biotin was removed using 7K Zeba columns for IA2, GAD, and TET (Thermo, Cat#89882). To ensure no loss of biotinylated INS, an Amicon 3kDa molecular weight cutoff centrifugal filter column was used to remove excess biotin and the top fraction was recovered (Millipore Sigma, Cat#UFC500396). Next, one mole of TotalSeqC PE Streptavidin was added to 4 moles of biotinylated antigen as follows: 0.5 ug of TotalSeqC PE Streptavidin was added to 1.54e-10 moles of biotinylated protein for 15 minutes on ice in the dark. This was repeated three times until a total of 2 ug (2 ug = 3.85e-11 moles) of TotalSeqC PE Streptavidin had been added. For samples 107, 108, 109, and 112, biotinylated insulin was combined with TotalSeq^TM^-C0961 PE Streptavidin (BioLegend, Cat#405155). All other samples were incubated with biotinylated insulin combined with TotalSeq^TM^-C0951 PE Streptavidin (BioLegend, Cat#405261). All samples were incubated with biotinylated tetramerized antigens as follows: GAD was combined with TotalSeq^TM^-C0954 PE Streptavidin (BioLegend, Cat#405267), IA2 was combined with TotalSeq^TM^-C0953 PE Streptavidin (BioLegend, Cat#405265), TET was combined with TotalSeq^TM^-C0952 PE Streptavidin (BioLegend, Cat#405263). For the “Empty” tetramer, TotalSeq^TM^-C0955 PE Streptavidin (BioLegend, Cat#405269) was quenched with biotin and excess biotin was removed using 7K Zebra columns (Thermo, Cat#89882). Validation of individual tetramers and appropriate dilutions to add to PBMCs was confirmed using flow cytometry. Tetramers were made fresh on the day of sample preparation for FACS sorting and 10X Genomics capture.

### Sample preparation for FACS sorting

Four cryopreserved PBMC samples were thawed and processed together, and eight samples were processed per day to minimize batch effects. Cells were thawed in a 37 C water bath, washed twice with 10 mL 1X PBS, and kept on ice for cell staining. Cells were stained with LIVE/DEAD^TM^ Fixable Near-IR Dead Cell Stain Kit at a 1:1500 dilution (ThermoFisher, Cat#L34975) in FACS buffer (1% BSA in 1X PBS) for 15 minutes on ice in the dark. Then, 2 ug of INS tetramer, 1 ug of IA2 and TET tetramers, and 0.75 ug of GAD and Empty tetramers in addition to 1:200 anti-human CD19-BV421 (BioLegend, Cat#363017) and 1:100 anti-human CD3-FITC (BioLegend, Cat#317305) were added to ∼30e6 PBMCs in 1 mL total volume. Cells were stained on ice for 20 minutes in the dark. Finally, cells were washed twice in FACS buffer (400 x g for 5 minutes) prior to analysis on the cell sorter.

### Sorting of PE-reactive B cells using FACS

After tetramer and antibody staining, cells were immediately brought to the CU Cancer Center Flow Cytometry Core. PE-binding B cells were sorted as Live/singlet/lymphocyte/CD19+/CD3-/PE+ cells into 100 uL of FACS buffer. Up to 20,000 B cells were collected, and in some instances, PE-negative binding B cells were also sorted to increase the overall yield of B cells to be captured.

### CITE-seq antibody staining

After the cells were sorted, they were stained with 1 uL each of the following TotalSeqC surface antibodies: anti-human IgM (BioLegend, Cat#314547), anti-human IgD (BioLegend, Cat#348245), anti-human CD27 (BioLegend, Cat#302853), anti-human CD21 (BioLegend, Cat#354923), and anti-human CXCR5 (BioLegend, Cat#356939). Cells were stained for 15 minutes on ice in the dark, washed once (400 x g for 5 minutes), then loaded onto the 10X Genomics system for cell capture at the CU Genomics Core. This data is unreported in the manuscript but present within the publicly available sequencing data.

### 5’ Single Cell Capture and Library Prep

The 10X Genomics Chromium system was used for 5’ mRNA capture with chip K according to user guide CG000330_ChromiumNextGEMSingleCell5-v2_CellSurfaceProtein_UserGuide_RevF.pdf. The deviations from this protocol included: 1.) Overloading the chip with up to 20,000 cells per well depending on the sample, and 2.) Using the Feature Barcode Kit Primer 4 (Cat# 2000277) to detect and amplify TotalSeq C barcodes.

Otherwise, the protocol was followed as written to produce gene expression, BCR V(D)J, and Feature Barcode Cell Surface Protein libraries for sequencing analysis. Library quality and quantity was determined using Agilent High Sensitivity D1000 TapeStation Screentape (Cat# 5067-5584), Reagents (Cat# 5067-5585), and Qubit DNA Quantification Assay (Cat# Q32851).

### Sequencing

The libraries produced for each sample were sequenced at 5,000 read pairs per cell for V(D)J and Cell Surface Protein Dual Index libraries and 20,000 read pairs per cell for the 5’ Gene Expression Dual Index library. They were sequenced paired end dual index on an Illumina NovaSeq6000 and demultiplexed all by the CU Genomics Core Staff.

### Bioinformatic Analysis

#### Pre-processing

Sequences from scRNA-seq were processed using 10x Genomics Cellranger v 7.1.0 software (*68*) using the human 10x genomics refdata-gex-GRCh38-2020-A reference downloaded from 10x genomics (https://www.10xgenomics.com/support/software/cell-ranger/downloads). V(D)J data was aligned using the 10x genomics reference refdata-cellranger-vdj-GRCh38-alts-ensembl-5.0.0 downloaded from 10x genomics (https://www.10xgenomics.com/support/software/cell-ranger/downloads). The antibody reference used can be found on our github https://github.com/CUAnschutzBDC/smith_nicholas_sc_antigen/blob/main/files/antibodies.csv. Raw data generated by Cellranger were then read into R v4.2.3 using the Seurat (*30*) v4.3.0 R package with at least 200 genes per cell and at least 3 cells. Cells were further filtered based on the number of genes per cell and the percent of mitochondrial reads per cell by finding outliers using ‘perCellQCMetrics’ and ‘perCellQCFilters’ from ‘scuttlè based on total RNA and ADT reads and features and percent mitochondria (*69*). The data were normalized by using ‘NormalizeDatà. For each sample, variable genes were found by using ‘FindVariableFeatures’ followed by removal of the immunoglobulin genes from the set and data was scaled using ‘ScaleDatà.

#### Doublet removal

Doublets were removed using ‘DoubletFinder’ (*70*) using the default values except for pK, nExp, and PCs. The pK was identified using the pK associated with the maximum BCmetric value after running ‘find.pK’ from doublet finder. All samples used PCs 1:20. After doublet removal, 2057 (110), 4240 (116), 4353 (108), 4466 (107), 2848 (113), 9058 (114), 8815 (118), 4911 (106), 7949 (117), 8497 (115), 9265 (105), 10547 (111), 9750 (102), 8609 (112), 11114 (119), 5135 (109) were used for downstream analysis.

#### Dimensionality reduction and clustering

Dimensionality reduction and clustering were performed using ‘RunPCÀ, ‘FindNeighbors’, ‘FindClusters’, and ‘RunUMAP’. ‘RunPCÀ was run using the default values except all variable features (minus the immunoglobulin genes) were used for the features argument. ‘FindNeighbors’ was run with default parameters except for the dims argument (different for each sample but ranging between 25 and 35). ‘FindClusters’ was run with default parameters except for the resolution argument (different for each sample but ranging between 0.6 and 1). ‘RunUMAP’ was run with default parameters except for the dims and metric arguments (the same number of dims as for ‘FindNeighbors’, metric=”correlation”).

#### Cell type identification

We named clusters using existing PBMC datasets (*30*) and (*15*) as references and determined cluster identity using ‘clustifyr’ (*71*). For each reference, we used our previously identified variable genes passed to ‘query_genes’ in the ‘clustify’ function. The top correlated cell type from any reference to each cluster was used as the cell type.

### Analysis of tetramers

#### Scar ambient background removal

Because we noticed high amounts of tetramer expression in empty droplets, we performed scar (*72*) (single cell ambient remover) on each sample individually. To run scar, we followed the recommended steps (https://scar-tutorials.readthedocs.io/en/latest/tutorials/scAR_tutorial_ambient_profile.html). First, we read in the raw and filtered ADT matrices. We then ran ‘scar.setup_anndatà to identify the ambient profile which we accessed with ‘filtered_object.uns[“ambient_profile_Antibody Capture”]’. Using this ambient profile and the raw data, we then ran ‘scar.model’, ‘scar.train’, and ‘scar.inferencè using the recommended arguments to pull out the ambient corrected ADT values.

#### Libra seq normalization

Libra seq normalization was performed on each sample individually as previously described (*27*). We read in the corrected tetramer matrices from running scar ambient background removal. We then used the following formula

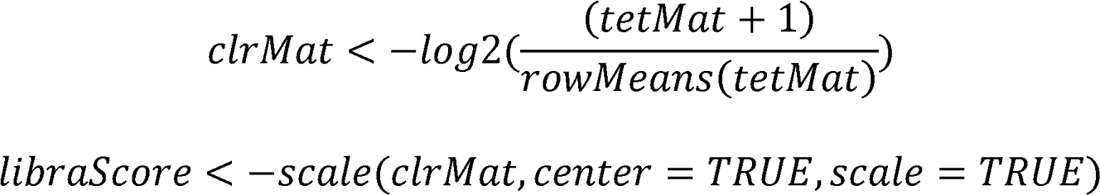

Any cells that had a value greater than 1 for the libra score for a tetramer were called positive for that tetramer.

#### Quantile normalization

Because we also had non-B cells captured in our sample, we used these to generate a second “quantile” tetramer score. Within each cell type individually, we used the cell types determined above and subset to only the non-B cells. We then read in the corrected tetramer matrices from running scar ambient background removal and identified the cutoff based on the 95^th^ quartile. We generated our score by dividing each tetramer count for each cell by the cutoff determined for the given tetramer. In this way, like the Libra score, any score greater than 1 is considered positive for a given tetramer while any score less than 1 is considered negative.

### Sample merging

#### Merging

The 16 samples were merged using the ‘mergè function from Seurat. The merged data was then normalized and scaled using ‘NormalizeDatà, ‘FindVariableFeatures’ followed by removal of the immunoglobulin genes, and ‘ScaleDatà as described above. Dimensionality reduction and clustering were performed using ‘RunPCÀ, ‘FindNeighbors’, ‘FindClusters’, and ‘RunUMAP’. ‘RunPCÀ was run using the default values except all variable features were used for the features argument (except for immunoglobulin genes). ‘FindNeighbors’ was run with default parameters except for the dims argument (dims=1:30). ‘FindClusters’ was run with default parameters except for the resolution argument (resolution=0.8). ‘RunUMAP’ was run with default parameters except for the dims and metric arguments (dims=1:30, metric=”correlation”).

#### Batch correction

We tested both ‘harmony’ (*73*) (using the default values except theta = 5) and ‘mnn’ (*31*) (using default values) batch correction methods on our data. After a thorough comparison between the two and exploration into how well cells that had been previously assigned as the same cell types overlapped, we chose to use the ‘mnn’ batch correction for all downstream analysis.

### Cell type identification

#### Clustifyr

We made a first pass at naming clusters existing PBMC datasets (*15, 30*) as references and determined cluster identity using ‘clustifyr’ (*71*). For each reference, we used our previously identified variable genes passed to ‘query_genes’ in the ‘clustify’ function. The top correlated cell type from any reference to each cluster was used as the first determination of cell type.

#### Cluster markers and pathways

To identify markers of each cluster, ‘FindAllMarkers’ was run using default settings except we set only.pos to TRUE. Genes were called differentially expressed if the adjusted p-value was less than 0.05 and the log fold change was greater than 0.5. We inspected these marker genes to determine if the cell types determined above were correct.

### V(D)J analysis

#### V(D)J statistics

V(D)J data was read in using ‘import_vdj’ from ‘djvdj’ (*74*) using the defaults except ‘filter_paired = FALSÈ and ‘include_mutations = TRUÈ. The values read in by ‘djvdj’ were used to determine V and J usage, CDR3 length, CDR3 charge, and percent somatic hypermutation. To calculate CDR3 charge, we used the ‘chargè function from ‘alakazam’ (*75*). Shannon diversity was calculated using ‘calc_diversity’ from ‘djvdj’ using the ‘shannon’ method from ‘abdiv’ (74, 82). Odds ratios and p-values were determined by finding the total count of each V gene for each comparison (T1D vs ND or AAB vs ND) and the total count of all cells for each comparison. The Odds ratio was calculated using the following formula

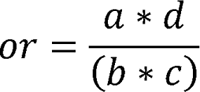

And the fishers exact test was run using

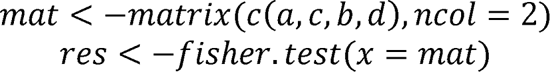

Where

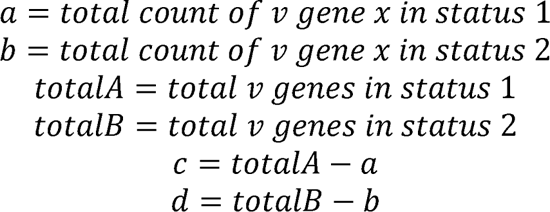

#### Clonal analysis

Clones were identified using the ‘AssignGenes.py’, ‘MakeDb.py’ and ‘DefineClones.py’ from the Change-O package in the immcantation suite (*75*). Clones were found both within and between samples. These clones were found based on CDR3 similarity using a hamming distance of 0.15 (85% similarity). We further refined these clones to have the same V and J call in addition to the CDR3 similarity calculated by ‘DefineClones.py’

### Differential expression

Differential expression analysis between each of the different status groups was performed using ‘FindMarkers’ from Seurat on each set of pairwise identities. Instead of using the default Wilcoxon test, we instead used the ‘MAST’ implementation. We performed DE using the same parameters on the twin sisters and the full family.

To find enriched pathways, we ran gProfiler2 (*76*) through the ‘run_gost’ function from the ‘scAnalysisR’ package (*77*, *78*). We then selected only the kegg pathways and clustered them using a custom function. This function first finds all genes associated with a particular keg term using ‘keggLink’ and ‘keggConv’ from the ‘KEGGREST’ (*79*) package and ‘mapIds’ from the ‘AnnotationDbì(*80*) package. Then, a matrix of jaccard distances for all pairwise gene sets is calculated. The jaccard distance was calculated with the following formula

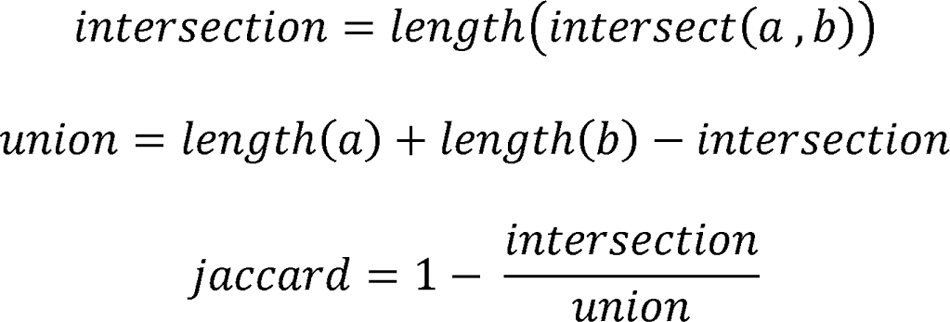

Where

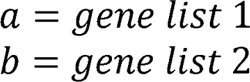

Finally, this matrix is converted into a distance matrix using the ‘as.dist’ function and clustered using ‘hclust’. We used this clustering to narrow down the kegg pathways into discrete groups as displayed.

### HLA typing

We identified HLA type using the ‘T1K’ (*81*) package. We performed HLA prediction using the example for 10x genomics data in their tutorial https://github.com/mourisl/T1K. Briefly, we used the R2 fastq as in input file, the R1 fastq as the barcode file, 0 15 + as the barcode range, the whitelist provided by 10x genomics as the barcodeWhitelist, and the hlaidx rna seq fasta file provided by the T1K package for the reference file. We used the output HLA alleles to determine the HLA types.

#### Recombinant monoclonal antibody production

Representative V(D)J sequences from AAB and T1D subjects identified by our quantile scoring approach that were part of an expanded IAR clone were chosen for antibody synthesis. B cells designated Islet_Multi reactive were chosen and validated for polyreactivity. Monoclonal antibodies were synthesized by Twist Bioscience. Briefly, sequences were cloned into the company’s human IgG1, human kappa chain, or human lambda chain expression vectors. Heavy and light chain genes were co-transfected into HEK293 cells. Secreted monoclonal antibodies were then purified from the supernatant using protein A agarose beads.

#### ELISA assay

High-protein binding microtiter plates (Corning, Cat#07-200-35) were coated overnight at 4 C with 10 ug/mL of antigen (INS, IA2, GAD, TET, DNA, or PE (Invitrogen, Cat#P801)) in 1X PBS. We included DNA of the same length and sequence as the TotalSeqC tags and PE as antigens to represent potential binding to the DNA barcode or PE fluorophore components of the tetramer probes. Plates were washed three times with 1X PBS 0.05% Tween and blocked with 1X PBS 2% BSA blocking buffer for one hour at room temperature. The plates were washed as before, and monoclonal antibodies were titrated down two-fold onto antigen coated wells from 40 – 0.625 ug/mL in triplicate. Antibodies were incubated for two hours at room temperature before washing. To detect antibody binding, goat anti human IgG (H+L) HRP (Invitrogen, Cat#31412) was added to wells at a 1:10,000 dilution for one hour at room temperature in the dark. Plates were washed and developed using TMB solution (Life Technologies, Cat#002023). The reaction was stopped with addition of 1N HCl and the plate was read at OD450nm using a BioTek Synergy H1 Microplate Reader. OD450 values from the blank wells included on each plate were subtracted to remove background signal from each test well prior to plotting with GraphPad Prism v9.

## Statistics

Statistical analysis was performed using R version 4.2.3. Comparisons between two groups were conducted using paired or unpaired Student’s t tests when comparing between samples given the small sample size. Bonferroni correction was always used when more than one comparison was run. Statistical analysis of sequencing data is described in depth above. Briefly, differential expression was run using MAST with multiple testing correction, gene set enrichment used a hypergeometric test, clustering of gene pathways was run based on jaccard distances and hierarchical clustering, differences in V gene usage was analyzed using an odds-ratio test, and any differences between status groups using all cells were determined using a Wilcoxon rank sum test.

## Author contributions

CAN, MJS, PAG, and KLW designed the research and developed the methodologies; FAT recruited, consented, and collected blood samples from subjects; HB was the study coordinator; CAN, MJS, SAE, and KPT performed experiments; CAN, MJS, and KLW analyzed the data; CAN, MJS and KLW prepared figures; CAN, MJS, and KLW wrote the original draft; all authors reviewed and edited the final draft; MJS and JRH provided funding.

## Data and materials availability

The data will be made available on GEO upon publication. All scripts and the analysis pipeline written in ‘snakemakè v6.3.0 to replicate this analysis will be made public on github (https://github.com/CUAnschutzBDC/smith_nicholas_sc_antigen) upon publication. All R analysis, scar, dropkick, and T1K were run through docker containers that are available on dockerhub (https://hub.docker.com/repositories/kwellswrasman-smith_2024_scar, smith_2024_dropkick, smith_2024_r_docker, smith_2024_t1k).

## Acknowledgments

We would like to thank the recent onset T1D patients, autoantibody positive participants, and nondiabetic first degree family members who were willing to donate blood for our study. We would also like to thank Liping Yu with the Barbara Davis Center Autoantibody Core for his help with validating the IA2 antigen and for providing GAD65 recombinant protein, Taylor Armstrong for her helpful discussions about rare HLA alleles and mapping them to common DR-DQ types, and Jenna Guthmiller for discussions regarding BCR repertoire analysis and recombinant antibody generation. Thank you to John Cambier for his careful review of this manuscript. Many thanks to the Barbara Davis Center Bioresource core facility for assistance with flow cytometry optimization, the CU Anschutz Cancer Center Flow Cytometry Shared Resource staff for assistance with cell sorting, and the CU Anschutz Genomics Core staff for 10X Genomics capture, library prep, and sequencing.

## Figures

Most figures were created using the ‘scAnalysisR’ package available on github (https://github.com/CUAnschutzBDC/scAnalysisR). Figure 1A was created using BioRender, Fig. 1B was created using FlowJo software v10.9, and Fig 7 was created using GraphPad Prism v9.

## Supplementary Materials

**Fig. S1.**
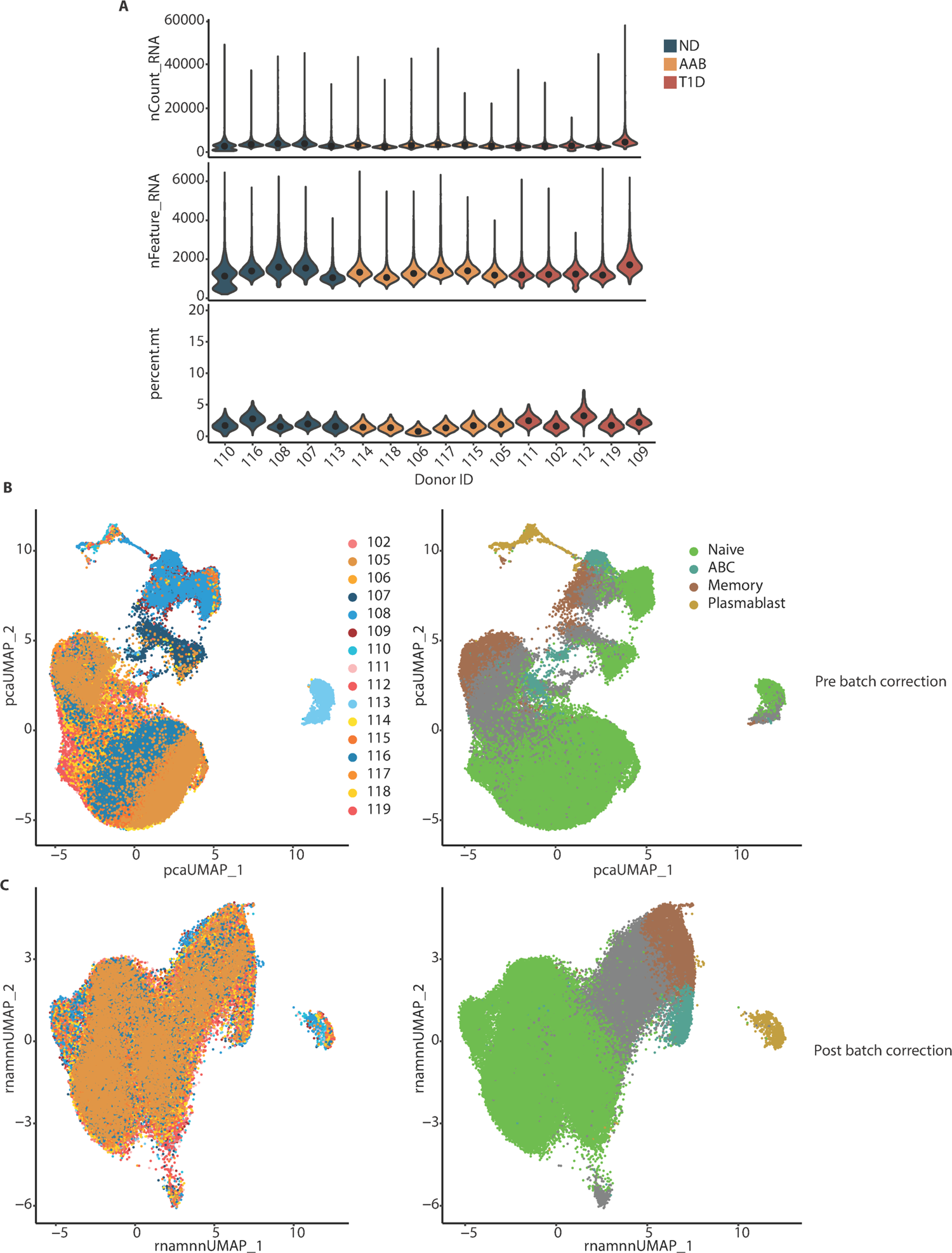
Quality metrics across single cell RNA-seq samples. **(A)** Violin plots of number of reads, number of genes and percent mitochondrial reads per cell post quality filtering across all 16 donors, colored by donor status group. **(B)** UMAP dimensionality reduction prior to batch correction, left colored by donor right colored by cell type. **(C)** UMAP dimensionality reduction following MNN batch correction, left colored by donor, right colored by cell type.

**Fig. S2.**
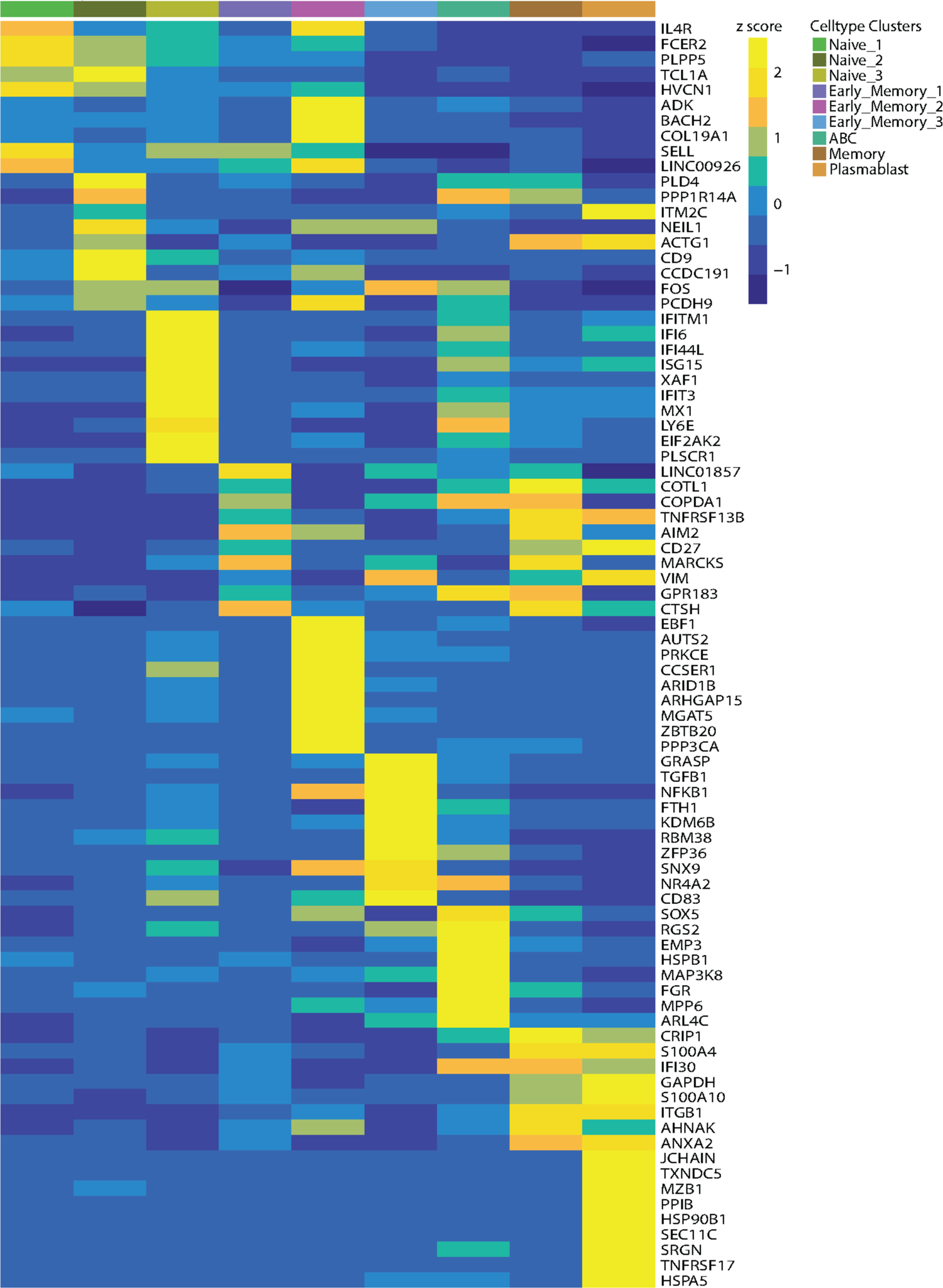
Top ten differentially expressed genes define nine unique B cell subsets. Heatmap of z-score of the top ten genes for each of the nine cell clusters across each cluster. Cluster names are indicated by color-coded labels across the top of the heatmap.

**Fig. S3.**
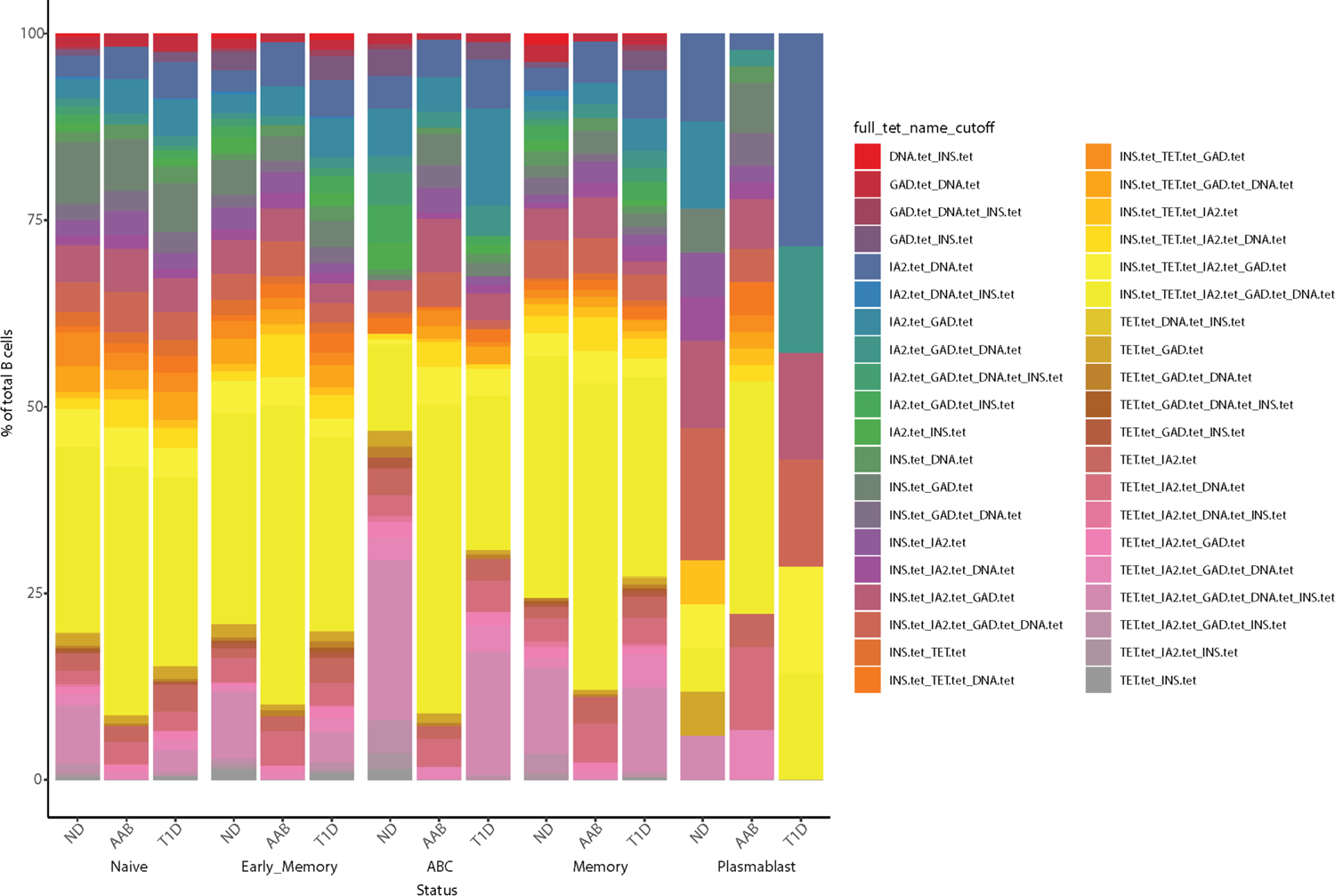
All antigen combinations occur across all B cells within the multi_islet reactivity group. Stacked bar plot divided by donor group and broad cell type showing all antigen combinations found across all B cells.

**Fig. S4.**
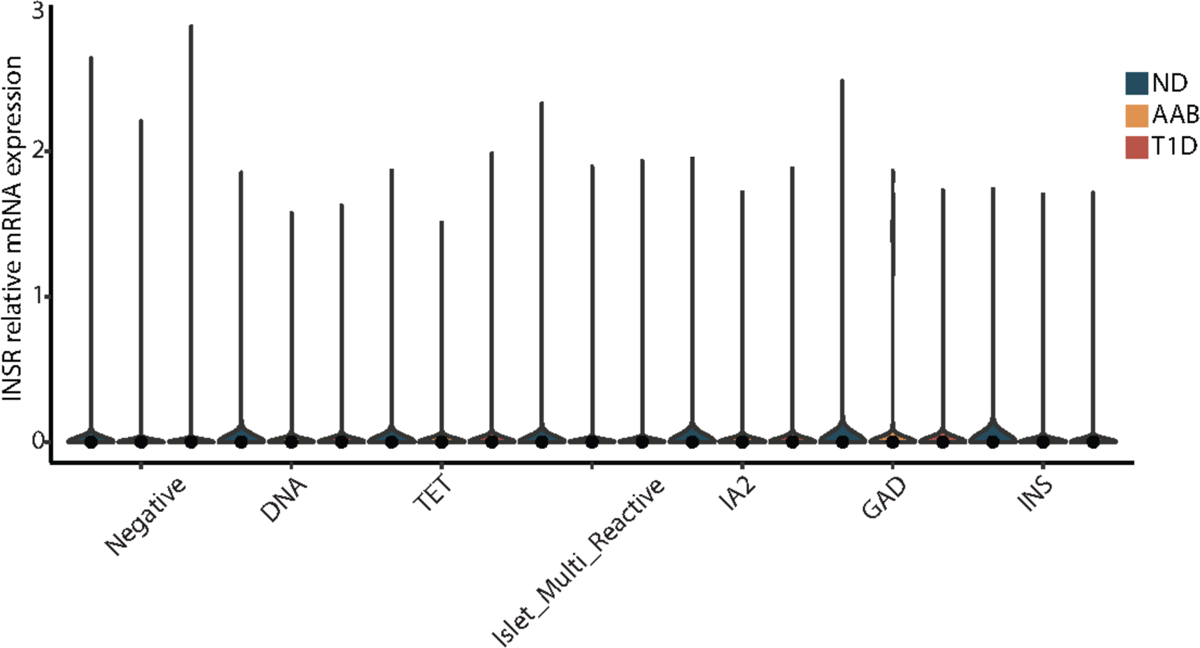
Insulin receptor expression is consistent across donor groups and BCR antigen reactivity types. Violin plot showing insulin receptor (INSR) mRNA gene expression levels for all antigen reactivities split by donor group.

**Fig. S5.**
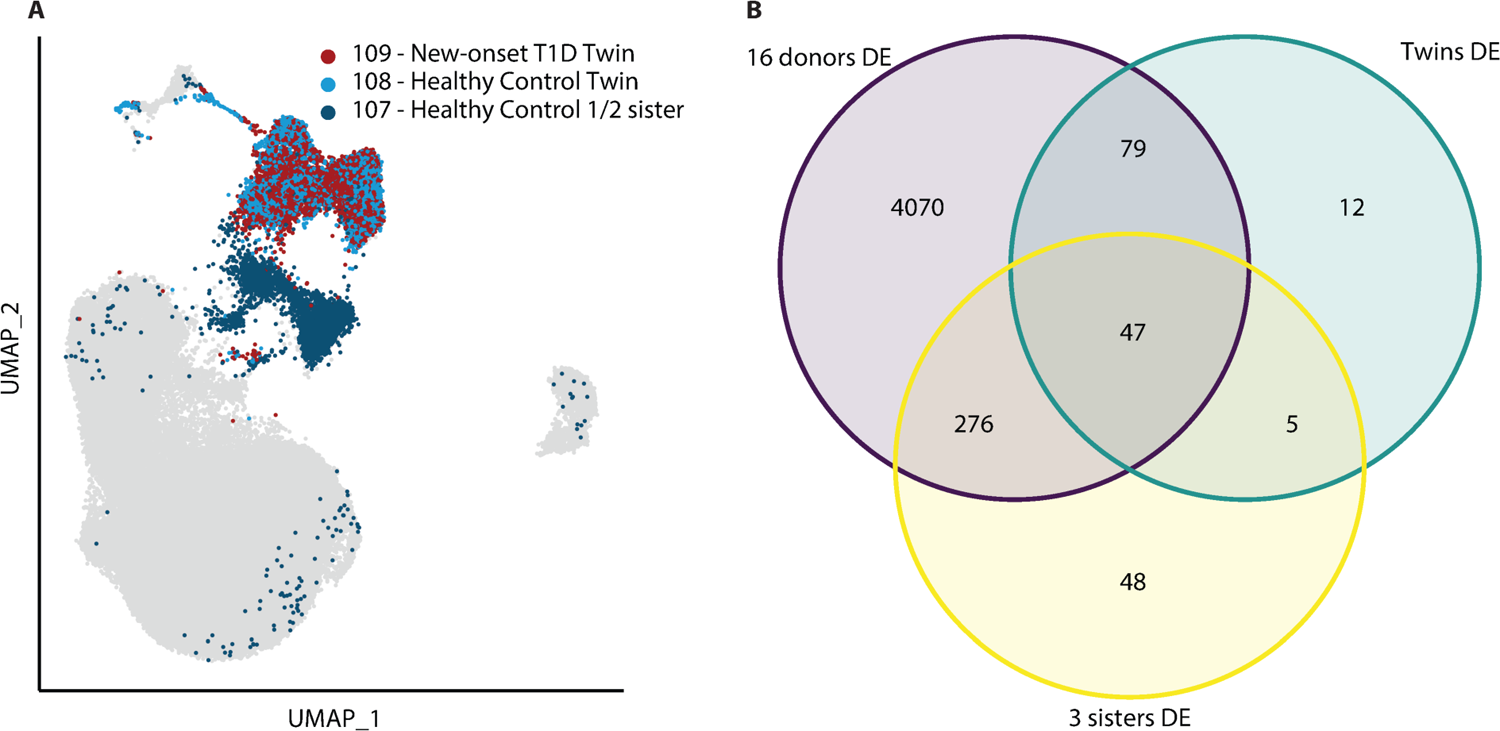
Differential expression between twin donors and their half-sister are consistent with differential expression between all donors. **(A)** Pre-batch corrected UMAP projections of all cells colored for sister-sibling donors 107, 108, and 109. **(B)** Venn diagram of differential gene expression for all donors (purple, top left), all three sisters (yellow, bottom), and the two twin sisters (blue, top right). Values within the sections of the venn diagram are the number of genes overlapping between comparisons.

**Fig. S6.**
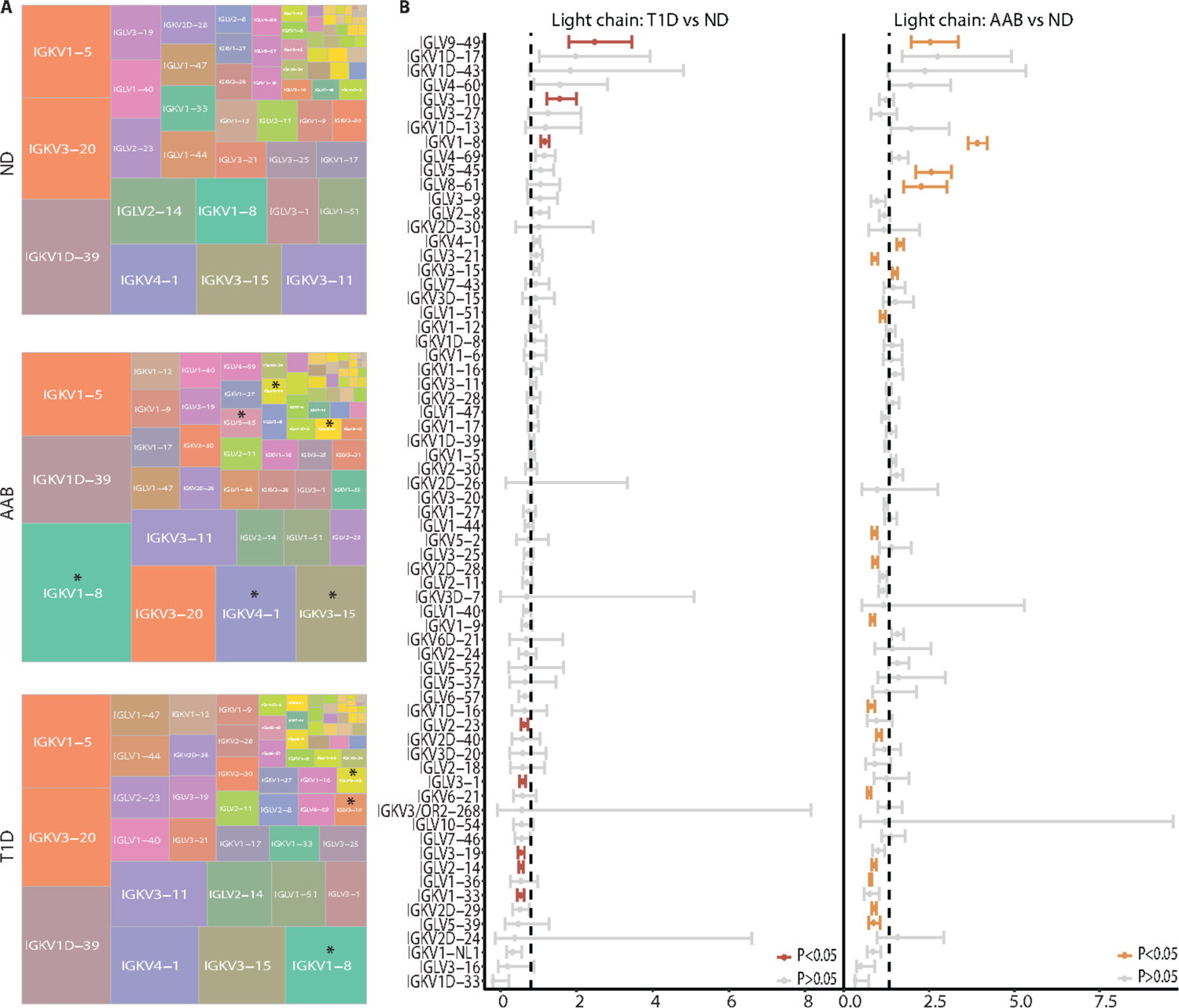
AAB and T1D donors have significant enrichment for particular light chain V gene usages. **(A)** Total B cells for each donor group were assessed for light chain V gene usage and plotted such that the size of each box is proportional to the frequency with which that gene was found among donors. Asterisks mark genes that were found to be significantly upregulated in AAB and T1D donors compared to ND. **(B)** Odds ratios for light chain V gene usage between T1D and ND donors and AAB and ND donors. P value is indicated by color where red and orange indicate p < 0.05 for T1D vs ND and AAB vs ND respectively.

**Fig. S7.**
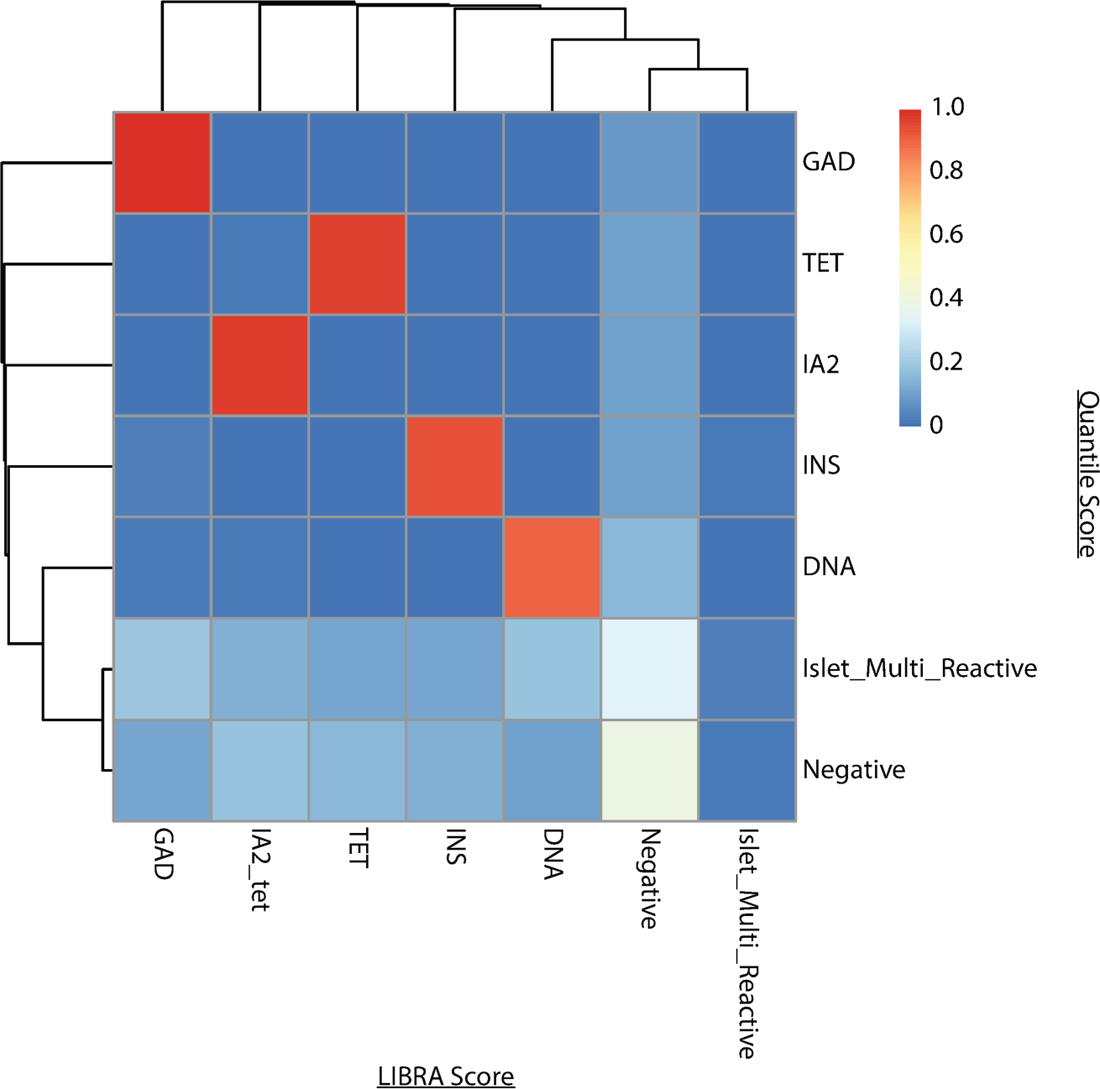
Comparison between the LIBRA-seq antigen reactivity calculation and our quantile normalization method. Heatmap of the score values assigned by either the Libra scoring method or quantile scoring method across antigen reactivity types.

## References

1. C. Hulbert, B. Riseili, M. Rojas, J. W. Thomas, B cell specificity contributes to the outcome of diabetes in nonobese diabetic mice. J. Immunol. 167, 5535–5538 (2001).

2. H. Noorchashm, Y. K. Lieu, N. Noorchashm, S. Y. Rostami, S. A. Greeley, A. Schlachterman, H. K. Song, L. E. Noto, A. M. Jevnikar, C. F. Barker, A. Naji, I-Ag7-mediated antigen presentation by B lymphocytes is critical in overcoming a checkpoint in T cell tolerance to islet beta cells of nonobese diabetic mice. J. Immunol. 163, 743–750 (1999).

3. E. Mariño, B. Tan, L. Binge, C. R. Mackay, S. T. Grey, B-cell cross-presentation of autologous antigen precipitates diabetes. Diabetes 61, 2893–2905 (2012).

4. D. V. Serreze, H. D. Chapman, D. S. Varnum, M. S. Hanson, P. C. Reifsnyder, S. D. Richard, S. A. Fleming, E. H. Leiter, L. D. Shultz, B lymphocytes are essential for the initiation of T cell-mediated autoimmune diabetes: analysis of a new “speed congenic” stock of NOD.Ig mu null mice. J. Exp. Med. 184, 2049–2053 (1996).

5. H. Noorchashm, S. A. Greeley, A. Naji, The role of t/b lymphocyte collaboration in the regulation of autoimmune and alloimmune responses. Immunol. Res. 27, 443–450 (2003).

6. J. Tian, D. Zekzer, L. Hanssen, Y. Lu, A. Olcott, D. L. Kaufman, Lipopolysaccharide-activated B cells down-regulate Th1 immunity and prevent autoimmune diabetes in nonobese diabetic mice. J. Immunol. 167, 1081–1089 (2001).

7. F. S. Wong, L. Wen, M. Tang, M. Ramanathan, I. Visintin, J. Daugherty, L. G. Hannum, C. A. Janeway Jr, M. J. Shlomchik, Investigation of the role of B-cells in type 1 diabetes in the NOD mouse. Diabetes 53, 2581–2587 (2004).

8. R. Mallone, V. Brezar, To B or not to B: (anti)bodies of evidence on the crime scene of type 1 diabetes?, Diabetes. 60 (2011)pp. 2020–2022.

9. D. G. Silva, S. R. Daley, J. Hogan, S. K. Lee, C. E. Teh, D. Y. Hu, K.-P. Lam, C. C. Goodnow, C. G. Vinuesa, Anti-islet autoantibodies trigger autoimmune diabetes in the presence of an increased frequency of islet-reactive CD4 T cells. Diabetes 60, 2102–2111 (2011).

10. M. D. Pescovitz, C. J. Greenbaum, H. Krause-Steinrauf, D. J. Becker, S. E. Gitelman, R. Goland, P. A. Gottlieb, J. B. Marks, P. F. McGee, A. M. Moran, P. Raskin, H. Rodriguez, D. A. Schatz, D. Wherrett, D. M. Wilson, J. M. Lachin, J. S. Skyler, Type 1 Diabetes TrialNet Anti-CD20 Study Group, Rituximab, B-lymphocyte depletion, and preservation of beta-cell function. N. Engl. J. Med. 361, 2143–2152 (2009).

11. K. C. Herold, M. D. Pescovitz, P. McGee, H. Krause-Steinrauf, L. M. Spain, K. Bourcier, A. Asare, Z. Liu, J. M. Lachin, H. M. Dosch, Type 1 Diabetes TrialNet Anti-CD20 Study Group, Increased T cell proliferative responses to islet antigens identify clinical responders to anti-CD20 monoclonal antibody (rituximab) therapy in type 1 diabetes. J. Immunol. 187, 1998–2005 (2011).

12. M. D. Pescovitz, C. J. Greenbaum, B. Bundy, D. J. Becker, S. E. Gitelman, R. Goland, P. A. Gottlieb, J. B. Marks, A. Moran, P. Raskin, H. Rodriguez, D. A. Schatz, D. K. Wherrett, D. M. Wilson, J. P. Krischer, J. S. Skyler, Type 1 Diabetes TrialNet Anti-CD20 Study Group, B-lymphocyte depletion with rituximab and β-cell function: two-year results. Diabetes Care 37, 453–459 (2014).

13. M. J. Smith, T. A. Packard, S. K. O’Neill, C. J. Henry Dunand, M. Huang, L. Fitzgerald-Miller, D. Stowell, R. M. Hinman, P. C. Wilson, P. A. Gottlieb, J. C. Cambier, Loss of anergic B cells in prediabetic and new-onset type 1 diabetic patients. Diabetes 64, 1703– 1712 (2015).

14. M. J. Smith, M. Rihanek, C. Wasserfall, C. E. Mathews, M. A. Atkinson, P. A. Gottlieb, J. C. Cambier, Loss of B-Cell Anergy in Type 1 Diabetes Is Associated With High-Risk HLA and Non-HLA Disease Susceptibility Alleles. Diabetes 67, 697–703 (2018).

15. Z. C. Stensland, C. A. Magera, H. Broncucia, B. D. Gomez, N. M. Rios-Guzman, K. L. Wells, C. A. Nicholas, M. Rihanek, M. J. Hunter, K. P. Toole, P. A. Gottlieb, M. J. Smith, Identification of an anergic BND cell-derived activated B cell population (BND2) in young-onset type 1 diabetes patients. J. Exp. Med. 220 (2023).

16. F. Pociot, Å. Lernmark, Genetic risk factors for type 1 diabetes. Lancet 387, 2331–2339 (2016).

17. J. M. Sosenko, J. S. Skyler, J. P. Palmer, J. P. Krischer, L. Yu, J. Mahon, C. A. Beam, D. C. Boulware, L. Rafkin, D. Schatz, G. Eisenbarth, Type 1 Diabetes TrialNet Study Group, Diabetes Prevention Trial-Type 1 Study Group, The prediction of type 1 diabetes by multiple autoantibody levels and their incorporation into an autoantibody risk score in relatives of type 1 diabetic patients. Diabetes Care 36, 2615–2620 (2013).

18. A. G. Ziegler, M. Rewers, O. Simell, T. Simell, J. Lempainen, A. Steck, C. Winkler, J. Ilonen, R. Veijola, M. Knip, E. Bonifacio, G. S. Eisenbarth, Seroconversion to multiple islet autoantibodies and risk of progression to diabetes in children. JAMA 309, 2473–2479 (2013).

19. P. A. Silveira, E. Johnson, H. D. Chapman, T. Bui, R. M. Tisch, D. V. Serreze, The preferential ability of B lymphocytes to act as diabetogenic APC in NOD mice depends on expression of self-antigen-specific immunoglobulin receptors. Eur. J. Immunol. 32, 3657– 3666 (2002).

20. D. V. Serreze, S. A. Fleming, H. D. Chapman, S. D. Richard, E. H. Leiter, R. M. Tisch, B lymphocytes are critical antigen-presenting cells for the initiation of T cell-mediated autoimmune diabetes in nonobese diabetic mice. J. Immunol. 161, 3912–3918 (1998).

21. A. Grandien, R. Fucs, A. Nobrega, J. Andersson, A. Coutinho, Negative selection of multireactive B cell clones in normal adult mice. Eur. J. Immunol. 24, 1345–1352 (1994).

22. R. Halverson, R. M. Torres, R. Pelanda, Receptor editing is the main mechanism of B cell tolerance toward membrane antigens. Nat. Immunol. 5, 645–650 (2004).

23. H. Wardemann, S. Yurasov, A. Schaefer, J. W. Young, E. Meffre, M. C. Nussenzweig, Predominant autoantibody production by early human B cell precursors. Science 301, 1374–1377 (2003).

24. R. Pelanda, R. M. Torres, Central B-cell tolerance: where selection begins. Cold Spring Harb. Perspect. Biol. 4, a007146 (2012).

25. S. B. Gauld, K. T. Merrell, J. C. Cambier, Silencing of autoreactive B cells by anergy: a fresh perspective. Curr. Opin. Immunol. 18, 292–297 (2006).

26. E. Meffre, A. Schaefer, H. Wardemann, P. Wilson, E. Davis, M. C. Nussenzweig, Surrogate light chain expressing human peripheral B cells produce self-reactive antibodies. J. Exp. Med. 199, 145–150 (2004).

27. I. Setliff, A. R. Shiakolas, K. A. Pilewski, A. A. Murji, R. E. Mapengo, K. Janowska, S. Richardson, C. Oosthuysen, N. Raju, L. Ronsard, M. Kanekiyo, J. S. Qin, K. J. Kramer, A. R. Greenplate, W. J. McDonnell, B. S. Graham, M. Connors, D. Lingwood, P. Acharya, L. Morris, I. S. Georgiev, High-Throughput Mapping of B Cell Receptor Sequences to Antigen Specificity. Cell 179, 1636–1646.e15 (2019).

28. M. Stoeckius, C. Hafemeister, W. Stephenson, B. Houck-Loomis, P. K. Chattopadhyay, H. Swerdlow, R. Satija, P. Smibert, Simultaneous epitope and transcriptome measurement in single cells. Nat. Methods 14, 865–868 (2017).

29. J. A. Noble, A. M. Valdes, M. Cook, W. Klitz, G. Thomson, H. A. Erlich, The role of HLA class II genes in insulin-dependent diabetes mellitus: molecular analysis of 180 Caucasian, multiplex families. Am. J. Hum. Genet. 59, 1134–1148 (1996).

30. Y. Hao, S. Hao, E. Andersen-Nissen, W. M. Mauck 3rd, S. Zheng, A. Butler, M. J. Lee, A. J. Wilk, C. Darby, M. Zager, P. Hoffman, M. Stoeckius, E. Papalexi, E. P. Mimitou, J. Jain, A. Srivastava, T. Stuart, L. M. Fleming, B. Yeung, A. J. Rogers, J. M. McElrath, C. A. Blish, R. Gottardo, P. Smibert, R. Satija, Integrated analysis of multimodal single-cell data. Cell 184, 3573–3587.e29 (2021).

31. L. Haghverdi, A. T. L. Lun, M. D. Morgan, J. C. Marioni, Batch effects in single-cell RNA-sequencing data are corrected by matching mutual nearest neighbors. Nat. Biotechnol. 36, 421–427 (2018).

32. I. C. Mouat, E. Goldberg, M. S. Horwitz, Age-associated B cells in autoimmune diseases. Cell. Mol. Life Sci. 79, 402 (2022).

33. H. W. King, N. Orban, J. C. Riches, A. J. Clear, G. Warnes, S. A. Teichmann, L. K. James, Single-cell analysis of human B cell maturation predicts how antibody class switching shapes selection dynamics. Sci Immunol 6 (2021).

34. A. Stewart, J. C.-F. Ng, G. Wallis, V. Tsioligka, F. Fraternali, D. K. Dunn-Walters, Single-Cell Transcriptomic Analyses Define Distinct Peripheral B Cell Subsets and Discrete Development Pathways. Front. Immunol. 12, 602539 (2021).

35. O. Franzén, L.-M. Gan, J. L. M. Björkegren, PanglaoDB: a web server for exploration of mouse and human single-cell RNA sequencing data. Database 2019 (2019).

36. M. Jourdan, A. Caraux, J. De Vos, G. Fiol, M. Larroque, C. Cognot, C. Bret, C. Duperray, D. Hose, B. Klein, An in vitro model of differentiation of memory B cells into plasmablasts and plasma cells including detailed phenotypic and molecular characterization. Blood 114, 5173–5181 (2009).

37. D. G. Efremov, S. Turkalj, L. Laurenti, Mechanisms of B Cell Receptor Activation and Responses to B Cell Receptor Inhibitors in B Cell Malignancies. Cancers 12 (2020).

38. A. Getahun, Role of inhibitory signaling in peripheral B cell tolerance. Immunol. Rev. 307, 27–42 (2022).

39. T. Tsubata, J. Wienands, B cell signaling. Introduction. Int. Rev. Immunol. 20, 675–678 (2001).

40. M. Perez-Andres, B. Paiva, W. G. Nieto, A. Caraux, A. Schmitz, J. Almeida, R. F. Vogt Jr, G. E. Marti, A. C. Rawstron, M. C. Van Zelm, J. J. M. Van Dongen, H. E. Johnsen, B. Klein, A. Orfao, Primary Health Care Group of Salamanca for the Study of MBL, Human peripheral blood B-cell compartments: a crossroad in B-cell traffic. Cytometry B Clin. Cytom. 78 **Suppl 1**, S47–60 (2010).

41. R. J. M. Bashford-Rogers, K. G. C. Smith, D. C. Thomas, Antibody repertoire analysis in polygenic autoimmune diseases. Immunology 155, 3–17 (2018).

42. S. Yurasov, H. Wardemann, J. Hammersen, M. Tsuiji, E. Meffre, V. Pascual, M. C. Nussenzweig, Defective B cell tolerance checkpoints in systemic lupus erythematosus. J. Exp. Med. 201, 703–711 (2005).

43. P. S. Linsley, F. Barahmand-Pour-Whitman, E. Balmas, H. A. DeBerg, K. J. Flynn, A. K. Hu, M. G. Rosasco, J. Chen, C. O’Rourke, E. Serti, V. H. Gersuk, K. Motwani, H. R. Seay, T. M. Brusko, W. W. Kwok, C. Speake, C. J. Greenbaum, G. T. Nepom, K. Cerosaletti, Autoreactive T cell receptors with shared germline-like α chains in type 1 diabetes. JCI Insight 6 (2021).

44. M. Nakayama, A. W. Michels, Using the T Cell Receptor as a Biomarker in Type 1 Diabetes. Front. Immunol. 12, 777788 (2021).

45. C. Lamagna, Y. Hu, A. L. DeFranco, C. A. Lowell, B cell-specific loss of Lyn kinase leads to autoimmunity. J. Immunol. 192, 919–928 (2014).

46. A. Getahun, N. A. Beavers, S. R. Larson, M. J. Shlomchik, J. C. Cambier, Continuous inhibitory signaling by both SHP-1 and SHIP-1 pathways is required to maintain unresponsiveness of anergic B cells. J. Exp. Med. 213, 751–769 (2016).

47. K. T. Coppieters, T. Boettler, M. von Herrath, Virus infections in type 1 diabetes. Cold Spring Harb. Perspect. Med. 2, a007682 (2012).

48. S. R. Isaacs, D. B. Foskett, A. J. Maxwell, E. J. Ward, C. L. Faulkner, J. Y. X. Luo, W. D. Rawlinson, M. E. Craig, K. W. Kim, Viruses and Type 1 Diabetes: From Enteroviruses to the Virome. Microorganisms 9 (2021).

49. M. J. Castleman, M. M. Stumpf, N. R. Therrien, M. J. Smith, K. E. Lesteberg, B. E. Palmer, J. P. Maloney, W. J. Janssen, K. J. Mould, J. D. Beckham, R. Pelanda, R. M. Torres, SARS-CoV-2 infection relaxes peripheral B cell tolerance. J. Exp. Med. 219 (2022).

50. P. Soulas, A. Woods, B. Jaulhac, A.-M. Knapp, J.-L. Pasquali, T. Martin, A.-S. Korganow, Autoantigen, innate immunity, and T cells cooperate to break B cell tolerance during bacterial infection. J. Clin. Invest. 115, 2257–2267 (2005).

51. K. E. Hansen, J. Arnason, A. J. Bridges, Autoantibodies and common viral illnesses. Semin. Arthritis Rheum. 27, 263–271 (1998).

52. E. Meffre, K. C. O’Connor, Impaired B-cell tolerance checkpoints promote the development of autoimmune diseases and pathogenic autoantibodies. Immunol. Rev. 292, 90–101 (2019).

53. M. B. Oldstone, Molecular mimicry and immune-mediated diseases. FASEB J. 12, 1255– 1265 (1998).

54. B. A. Suliman, Potential clinical implications of molecular mimicry-induced autoimmunity. Immun Inflamm Dis 12, e1178 (2024).

55. B. Sundaresan, F. Shirafkan, K. Ripperger, K. Rattay, The Role of Viral Infections in the Onset of Autoimmune Diseases. Viruses 15 (2023).

56. N. H. Trier, G. Houen, Antibody Cross-Reactivity in Auto-Immune Diseases. Int. J. Mol. Sci. 24 (2023).

57. Y. A. Lomakin, M. Y. Zakharova, A. V. Stepanov, M. A. Dronina, I. V. Smirnov, T. V. Bobik, A. Y. Pyrkov, N. V. Tikunova, S. N. Sharanova, V. M. Boitsov, S. Y. Vyazmin, M. R. Kabilov, A. E. Tupikin, A. N. Krasnov, N. A. Bykova, Y. A. Medvedeva, M. V. Fridman, A. V. Favorov, N. A. Ponomarenko, M. V. Dubina, A. N. Boyko, V. V. Vlassov, A. A. Belogurov Jr, A. G. Gabibov, Heavy–light chain interrelations of MS-associated immunoglobulins probed by deep sequencing and rational variation. Mol. Immunol. 62, 305–314 (2014).

58. M. E. Chriswell, A. R. Lefferts, M. R. Clay, A. R. Hsu, J. Seifert, M. L. Feser, C. Rims, M. S. Bloom, E. A. Bemis, S. Liu, M. D. Maerz, D. N. Frank, M. K. Demoruelle, K. D. Deane, E. A. James, J. H. Buckner, W. H. Robinson, V. M. Holers, K. A. Kuhn, Clonal IgA and IgG autoantibodies from individuals at risk for rheumatoid arthritis identify an arthritogenic strain of Subdoligranulum. Sci. Transl. Med. 14, eabn5166 (2022).

59. J. Vencovský, E. Zd’árský, S. P. Moyes, A. Hajeer, Š. Ruzicková, Z. Cimburek, W. E. Ollier, R. N. Maini, R. A. Mageed, Polymorphism in the immunoglobulin VH gene V1 69 affects susceptibility to rheumatoid arthritis in subjects lacking the HLA DRB1 shared epitope. Rheumatology 41, 401–410 (2002).

60. J. C. W. Edwards, L. Szczepanski, J. Szechinski, A. Filipowicz-Sosnowska, P. Emery, D. R. Close, R. M. Stevens, T. Shaw, Efficacy of B-cell-targeted therapy with rituximab in patients with rheumatoid arthritis. N. Engl. J. Med. 350, 2572–2581 (2004).

61. S. E. Baranzini, M. C. Jeong, C. Butunoi, R. S. Murray, C. C. Bernard, J. R. Oksenberg, B cell repertoire diversity and clonal expansion in multiple sclerosis brain lesions. J. Immunol. 163, 5133–5144 (1999).

62. I. Mikocziova, V. Greiff, L. M. Sollid, Immunoglobulin germline gene variation and its impact on human disease. Genes Immun. 22, 205–217 (2021).

63. H. Yuuki, T. Itamiya, Y. Nagafuchi, M. Ota, K. Fujio, B cell receptor repertoire abnormalities in autoimmune disease. Front. Immunol. 15, 1326823 (2024).

64. M. J. Smith, J. C. Cambier, P. A. Gottlieb, Endotypes in T1D: B lymphocytes and early onset. Curr. Opin. Endocrinol. Diabetes Obes. 27, 225–230 (2020).

65. R. M. Hinman, J. C. Cambier, Role of B lymphocytes in the pathogenesis of type 1 diabetes. Curr. Diab. Rep. 14, 543 (2014).

66. M. J. Redondo, N. G. Morgan, Heterogeneity and endotypes in type 1 diabetes mellitus. Nat. Rev. Endocrinol. 19, 542–554 (2023).

67. T. Amendt, P. Yu, TLR7 and IgM: Dangerous Partners in Autoimmunity. Antibodies (Basel) 12 (2023).

68. G. X. Y. Zheng, J. M. Terry, P. Belgrader, P. Ryvkin, Z. W. Bent, R. Wilson, S. B. Ziraldo, T. D. Wheeler, G. P. McDermott, J. Zhu, M. T. Gregory, J. Shuga, L. Montesclaros, J. G. Underwood, D. A. Masquelier, S. Y. Nishimura, M. Schnall-Levin, P. W. Wyatt, C. M. Hindson, R. Bharadwaj, A. Wong, K. D. Ness, L. W. Beppu, H. J. Deeg, C. McFarland, K. R. Loeb, W. J. Valente, N. G. Ericson, E. A. Stevens, J. P. Radich, T. S. Mikkelsen, B. J. Hindson, J. H. Bielas, Massively parallel digital transcriptional profiling of single cells. Nat. Commun. 8, 14049 (2017).

69. D. J. McCarthy, K. R. Campbell, A. T. L. Lun, Q. F. Wills, Scater: pre-processing, quality control, normalization and visualization of single-cell RNA-seq data in R. Bioinformatics 33, 1179–1186 (2017).

70. C. S. McGinnis, L. M. Murrow, Z. J. Gartner, DoubletFinder: Doublet Detection in Single-Cell RNA Sequencing Data Using Artificial Nearest Neighbors. Cell Syst 8, 329–337.e4 (2019).

71. R. Fu, A. E. Gillen, R. M. Sheridan, C. Tian, M. Daya, Y. Hao, J. R. Hesselberth, K. A. Riemondy, clustifyr: an R package for automated single-cell RNA sequencing cluster classification. F1000Res. 9, 223 (2020).

72. C. Sheng, R. Lopes, G. Li, S. Schuierer, A. Waldt, R. Cuttat, S. Dimitrieva, A. Kauffmann, E. Durand, G. G. Galli, G. Roma, A. de Weck, Probabilistic machine learning ensures accurate ambient denoising in droplet-based single-cell omics, bioRxiv (2022). 10.1101/2022.01.14.476312.

73. I. Korsunsky, N. Millard, J. Fan, K. Slowikowski, F. Zhang, K. Wei, Y. Baglaenko, M. Brenner, P.-R. Loh, S. Raychaudhuri, Fast, sensitive and accurate integration of single-cell data with Harmony. Nat. Methods 16, 1289–1296 (2019).

74. Djvdj: An R Package to Analyze Single-Cell V(D)J Data (Github; https://github.com/rnabioco/djvdj).

75. N. T. Gupta, J. A. Vander Heiden, M. Uduman, D. Gadala-Maria, G. Yaari, S. H. Kleinstein, Change-O: a toolkit for analyzing large-scale B cell immunoglobulin repertoire sequencing data. Bioinformatics 31, 3356–3358 (2015).

76. L. Kolberg, U. Raudvere, I. Kuzmin, J. Vilo, H. Peterson, gprofiler2 -- an R package for gene list functional enrichment analysis and namespace conversion toolset g:Profiler. F1000Res. 9 (2020).

77. scAnalysisR: A Package to Analyze Single Cell Data in R. This Package Is under Active Development (Github; https://github.com/CUAnschutzBDC/scAnalysisR).

78. K. L. Wells, C. N. Miller, A. R. Gschwind, W. Wei, J. D. Phipps, M. S. Anderson, L. M. Steinmetz, Combined transient ablation and single-cell RNA-sequencing reveals the development of medullary thymic epithelial cells. Elife 9 (2020).

79. Tenenbaum D, Maintainer B (2024). KEGGREST: Client-side REST access to the Kyoto Encyclopedia of Genes and Genomes (KEGG). R package version 1.44.0, https://bioconductor.org/packages/KEGGREST.

80. Pagès H, Carlson M, Falcon S, Li N (2024). AnnotationDbi: Manipulation of SQLite-based annotations in Bioconductor. R package version 1.66.0, https://bioconductor.org/packages/AnnotationDbi.

81. L. Song, G. Bai, X. S. Liu, B. Li, H. Li, Efficient and accurate KIR and HLA genotyping with massively parallel sequencing data. Genome Res. 33, 923–931 (2023).

82. abdiv: Alpha and Beta Diversity Measures.(CRAN; https://CRAN.R-project.org/package=abdiv)

