## Supplementary_File_3 for "Islet-antigen reactive B cells display a unique phenotype and BCR repertoire in autoantibody positive and recent-onset type 1 diabetes patients"

**File 3. IA2 sequence used for protein expression.** Exact sequence synthesized by IDT to express IA2 for use in antigen tetramer reagents.

cgccagcaggataaagaacgcctggcggcgctgggcccggaaggcgcgcatggcgataccacctttgaatatcaggatctgtgccgccagcatatggcgaccaaaagcctgtttaaccgcgcggaaggcccgccggaaccgagccgcgtgagcagcgtgagcagccagtttagcgatgcggcgcaggcgagcccgagcagccatagcagcaccccgagctggtgcgaagaaccggcgcaggcgaacatggatattagcaccggccatatgattctggcgtatatggaagatcatctgcgcaaccgcgatcgcctggcgaaagaatggcaggcgctgtgcgcgtatcaggcggaaccgaacacctgcgcgaccgcgcagggcgaaggcaacattaaaaaaaaccgccatccggattttctgccgtatgatcatgcgcgcattaaactgaaagtggaaagcagcccgagccgcagcgattatattaacgcgagcccgattattgaacatgatccgcgcatgccggcgtatattgcgacccagggcccgctgagccataccattgcggatttttggcagatggtgtgggaaagcggctgcaccgtgattgtgatgctgaccccgctggtggaagatggcgtgaaacagtgcgatcgctattggccggatgaaggcgcgagcctgtatcatgtgtatgaagtgaacctggtgagcgaacatatttggtgcgaagattttctggtgcgcagcttttatctgaaaaacgtgcagacccaggaaacccgcaccctgacccagtttcattttctgagctggccggcggaaggcaccccggcgagcacccgcccgctgctggattttcgccgcaaagtgaacaaatgctatcgcggccgcagctgcccgattattgtgcattgcagcgatggcgcgggccgcaccggcacctatattctgattgatatggtgctgaaccgcatggcgaaaggcgtgaaagaaattgatattgcggcgaccctggaacatgtgcgcgatcagcgcccgggcctggtgcgcagcaaagatcagtttgaatttgcgctgaccgcggtggcggaagaagtgaacgcgattctgaaagcgctgccgcag
